# Active Cell Divisions Generate Exotic Fourfold Orientationally Ordered Phase in Living Tissue

**DOI:** 10.1101/2021.07.28.453899

**Authors:** Dillon Cislo, Haodong Qin, Fengshuo Yang, Mark J. Bowick, Sebastian J. Streichan

## Abstract

Morphogenesis, the process through which genes generate form, establishes tissue scale order as a template for constructing the complex shapes of the body plan. The extensive growth required to build these ordered substrates is fuelled by cell proliferation, which, naively, should destroy order. The active mechanisms that couple cellular and physical processes to generate and maintain global order, thereby reconciling this seeming contradiction, remain elusive. Using live imaging and tissue cartography, we quantitatively analyze the dynamics of fourfold tissue ordering in the crustacean *Parhyale hawaiensis*. We show that cell divisions are the main drivers of tissue flow leading to a fourfold orientationally ordered phase. Waves of anisotropic cell proliferation propagate across the embryo with precise choreography, such that defects introduced into the nascent lattice by cell divisions are healed by subsequent divisions through active defect climb. Orchestrating cell proliferation rates and orientations enables cell divisions to organize, rather than fluidize, the tissue. The result is a robust, active mechanism for generating global orientational order in a non-equilibrium system that sets the stage for the subsequent development of shape and form.

---

Order is necessary within living tissues to serve as a substrate for complex structure. The crucial role that order plays in morphogenesis is particularly apparent in direct developers. These animals assemble a complete, miniature version of the adult body during embryogenesis [1]. The limbs and organs comprising the adult form are arranged according to a specific body plan that ensures proper biomechanical functionality [2, 3]. This body plan is symmetry broken. Body parts are aligned and oriented relative to distinct principal body axes [3]. In order to reliably generate the correct arrangement of limbs and organs, direct developers create organizational templates from ordered regions of tissue, akin to a coordinate system spanning the entire body.

Such templates must be ordered to delineate the body plan, but also retain sufficient fluidity to facilitate the large deformations typical of development. Orientational order, an intermediate state between solid and and liquid matter, has been previously studied in non-living, thermally equilibrated systems [4–8]. More recently, orientational order has been demonstrated in the late stages of development, where organs use planar polarized signals to arrange cells into an ordered phase in the absence of proliferation [9–11]. In contrast, the initial structuring of the body plan during early embryonic stages occurs via the sequential outgrowth of segments and is fuelled by cell proliferation [1]. Generically, cell proliferation should fluidize any tissue [12, 13]. Fluidity gives rise to rearrangements that mix cells and drive the tissue away from an ordered state [13, 14]. Orientationally ordered phases occupy only small fractions of their respective phase spaces in thermally equilibrated systems [15–17]. Thus, it remains unclear how non-equilibrium mechanisms in living systems can generate the requisite order to specify the body plan in the presence of cell divisions.

Here, we used *Parhyale hawaiensis*, an emerging model system of direct limb morphogenesis [18, 19], to study the interplay of growth and order. *Parhyale* implements its body plan sequentially via extensive cell proliferation [20]. Prior to appendage outgrowth, the ectoderm forms a grid of locally ordered cells [20, 21] (Fig. 1, A and C to E), a feature shared among malacostracans [19]. The rows of this grid correspond to segments of the adult body [20, 21]. Limb buds form at specific locations in the grid and give rise to numerous functionally specific appendages [18]. Importantly, limb orientation, in terms of the dorso-ventral (D-V) and antero-posterior axes (A-P), can be traced back to the local arrangement of precursor cells at the grid stage [20, 22].

**Fig. 1.**
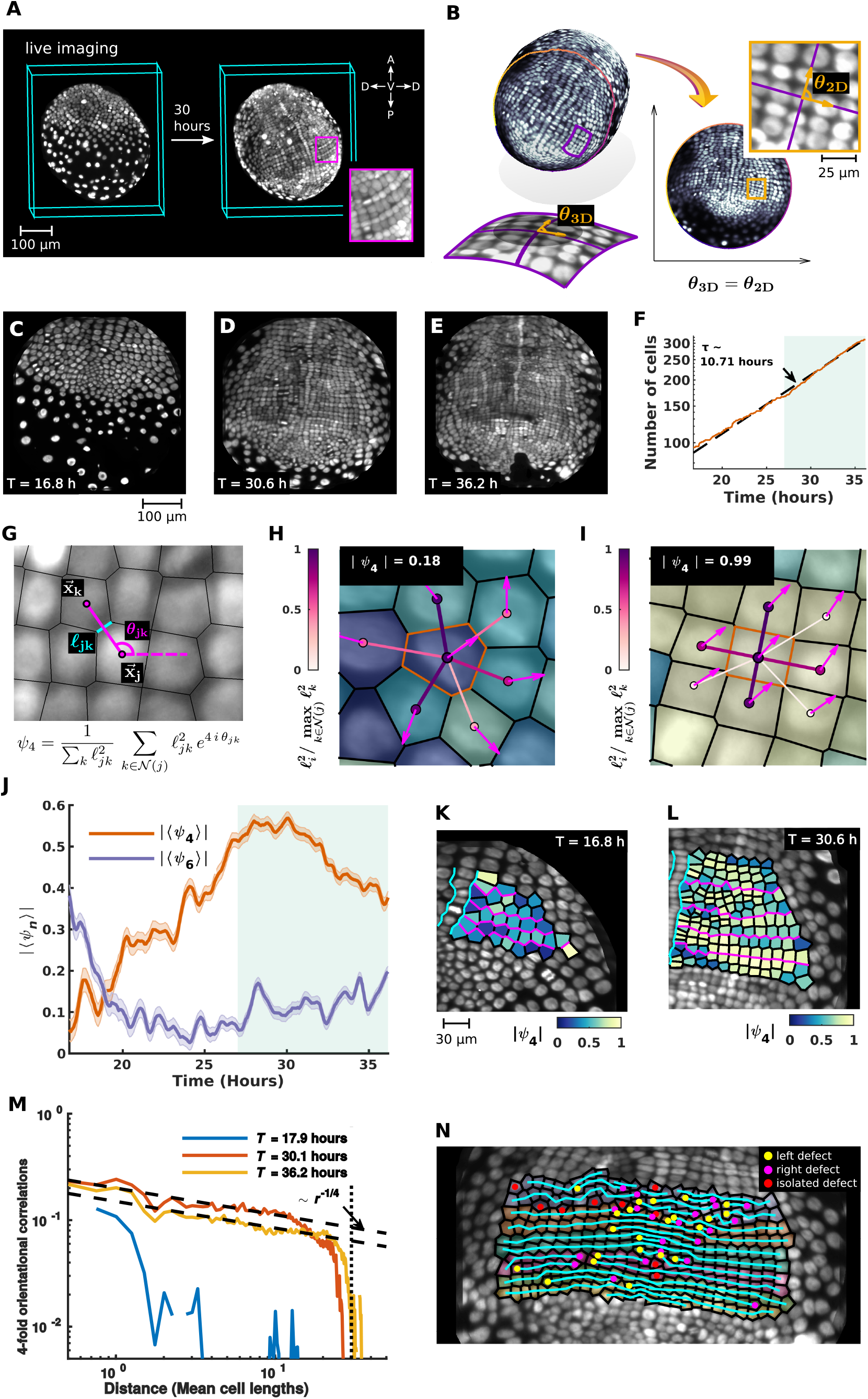
Live imaging reveals the dynamics of a fourfold orientationally ordered phase in living tissue. (**A**) Multiview light sheet microscopy allows for non-invasive 3D imaging of growing *Parhyale hawaiensis* embryos. (**B**) Conformal tissue cartography faithfully captures relative orientations of cells and streamlines data analysis via dimensional reduction. (**C** to **E**) The trunk ectodermal germband pulled back to the plane by tissue cartography. (**F**) The number of cells in the tissue, shown on a log scale. Dotted line is an exponential fit to the cell doubling time. (**G)** A schematic of the discrete *n*-fold complex orientational order parameter. Here, 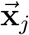 and 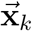 denote the centroids of cells *j* and *k*, respectively, *θ*_*jk*_ denotes the angle between the horizontal axis and the separation vector 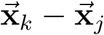, and *ℓ*_*jk*_ denotes the length of the Voronoi edge shared by cells *j* and *k*. The sum runs over all cells *k* in the neighborhood of cell *j*, i.e. *k ∈* 𝒩 (*j*). The weighting of the sum by the squared Voronoi edge lengths 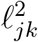 reduces the influence of noise in centroid positions on the final value of the orientational order parameter. (**H** and **I**) Specific examples of the construction of the single cell order parameter. Value shown in the inset is the magnitude of the order parameter of the seed cell highlighted by an orange boundary. Cell color and arrows indicate the magnitude and phase of the single cell order parameters, respectively. The color and thickness of the bonds between the seed cell and its neighbors indicate the relative weight with which each bond contributes to the sum defining the order parameter. (H) A disordered cell. (I) A highly ordered cell. (**J)** The absolute value of the mean fourfold and sixfold orientational order parameters over the whole embryo. Shaded contours show standard errors. (**K** and **L**) The absolute value of the single cell fourfold orientational order parameter in the left ectodermal compartment. (**M**) The two-point correlator of the fourfold orientational order parameter. Vertical line shows the largest lateral dimension of the system. (**N**) Edge defects introduced into rows of cells by division.

We performed whole embryo live imaging of *Parhyale* using multiview light sheet microscopy [23]. Transgenic embryos with fluorescently labeled nuclei were imaged for 35 hours beginning three days post fertilization (Fig. 1A). For the duration of this period, the ectoderm is a monolayer [20] and can be well approximated as a curved 2D surface. Tissue cartography could therefore be used to facilitate data analysis by exploiting the inherent low dimensionality of the system [24]. The surface of interest corresponding to the ectoderm was dynamically extracted at each time point via a combination of machine learning [25] and level-set methods [26] (see supplementary text). We then used discrete Ricci flow [27] to define conformal charts that mapped the curved surface into the plane in a way that preserved angles (Fig. 1, B to E). Conformal tissue cartography ensures that the relative cell orientations, and therefore the orientational order, can be faithfully quantified from the planar data.

Working in the planar pullback space, we implemented automated image segmentation routines to detect cells (see supplementary text). We found that the number of cells increases exponentially with a typical doubling time of 10 h (Fig. 1F). Next, we constructed a complex order parameter to quantify the relative orientations of neighboring cells (Fig. 1G). Cell position was taken to be the center of mass of the nuclei. Instantaneous cell-cell connectivity was approximated by Voronoi tessellation. The order parameter assigns to each cell a pair of quantitative measures, magnitude and phase, indicating the extent to which neighboring cells are coherently positioned according to a specific *n*-fold lattice structure (fig. S1) and the local orientation of the ordered neighborhood, respectively. The magnitude ranges between 0 (no *n*-fold order) and 1 (maximal *n*-fold order). For *n* = 4, the magnitude of the order parameter peaks when all neighbors of a cell are organized in a rectangular fashion (Fig. 1, H and I). Our analysis reveals that the tissue initially exhibits no four fold order (Fig. 1, J and K). Gradually, as the number of cells grows, the tissue adopts an increasingly fourfold ordered state, which peaks in magnitude around 30 h (Fig. 1, J and L). No significant six fold order was detected at any time during this stage (Fig. 1J and fig. S2A).

We also calculated the two point correlation functions of the order parameter, which measure the agreement of the magnitude and phase of local order between cells as a function of their separation (Fig. 1M and fig. S2A). At early times, orientational order is short range, restricted to less than a cell length. In contrast, at later times, when the global fourfold order parameter peaks, orientational order is quasi-long range, with correlations that decay algebraically across the entire surface (Fig. 1M). Therefore, strongly ordered local cell neighborhoods are coherently ordered, in both magnitude and phase, across the whole embryo.

Next, we tested if the ectodermal grid exhibits translational order, which would be reflected in a periodic positioning of cells along the D-V or A-P axis (fig. S4). Instead, analysis of the pairwise radial distribution function indicates cell spacing is not periodic along any axis (fig. S2, B and C). Thus, the tissue exhibits neither translational nor smectic order, despite its seeming regularity. The significance of these measurements was confirmed by extensive validation against synthetic data sets with similar cell densities and system sizes (fig. S3 and supplementary text). This lack of periodic cell spacing can be partly explained by the presence of edge defects in rows of cells. These defects are disordered, reminiscent of the defect gases characteristic of orientationally ordered phases (Fig. 1N). While we observe only a few isolated defects, many defects can be associated with one another by a row of ordered cells with a defect on the left and right sides, respectively (Fig. 1N). Together, these results show that the tissue achieves a true orientationally ordered phase extending over the entire trunk ectodermal germband.

We performed single cell tracking to reconstruct the flow fields that organize the ectoderm during the rise of fourfold order. A parasegment pre-cursor row (PSPR) is the fundamental supercellular unit of morphogenesis in the trunk ectodermal germband during this phase of growth [20, 21]. A PSPR is a single row of cells, oriented perpendicularly to the A-P axis, that can be directly associated with particular segments of the adult body. As a unit, PSPRs are appended to the grid sequentially. Cells are recruited from a pool of unorganized ectoderm at the posterior pole of the embryo and assembled into nascent rows so that each newly constructed PSPR lies immediately posterior to the PSPR that preceded it (Fig. 2A). Despite being arranged as a row, newly built PSPRs do not yet exhibit fourfold order (Fig. 1J and fig. S6).

**Fig. 2.**
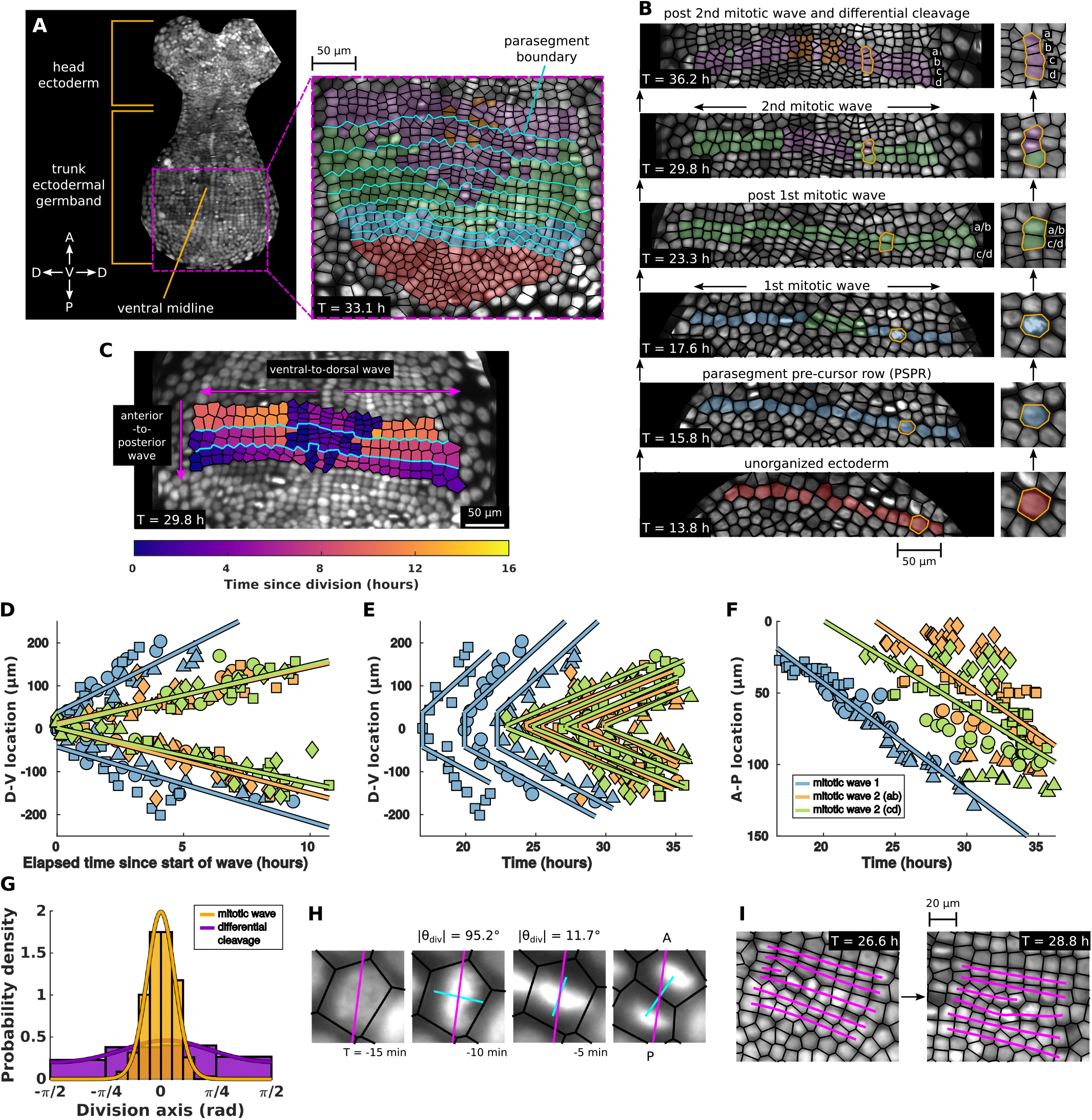
Waves of actively oriented divisions generate fourfold order. (**A** and **B**) A schematic of parasegment formation and cell proliferation in the trunk ectodermal germband. Inset in (A) shows the spatial distribution of cell populations delineated by number of divisions since parasegment formation once parasegments are assembled from the pool of unorganized ectoderm at the posterior pole. (B) shows the proliferation of a single parasegment. (**C**) Snapshot of elapsed time since division reveals two orthogonal phase waves within and across parasegments. (**D** to **F**) The location of mitotic wave division events over time. Shapes indicate the parasegment within which a division occurs. Indicated lines are linear fits to all division events associated to a particular mitotic wave. (D) and **(E)** show the location of each division along the D-V axis. Division times in (D) have been normalized to the occurrence of the first division event associated with a particular wave in a specific parasegment. The speed of mitotic wave 1 is 19.2 ± 2.1 *µ*m/hr. The speed of mitotic wave 2 (AB) is 13.9 ± 1.2 *µ*m/hr. The speed of mitotic wave 2 (CD) is 13.1 ± 0.9 *µ*m/hr. (F) shows the location of each division along the A-P axis. The speed of mitotic wave 1 is 7.5 ± 0.3 *µ*m/hr. The speed of mitotic wave 2 (AB) is 6.9 ± 0.9 *µ*m/hr. The speed of mitotic wave 2 (CD) is 6.1 ± 0.9 *µ*m/hr. (**G**) The orientation of cell division axes relative to the A-P axis. Indicated curves are von Mises distributions fit to histogram counts. The circular mean division angle and angular deviation for the mitotic waves are 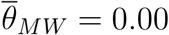 rad and *s*_*MW*_ = 0.41. The circular mean division angle and angular deviation for the differential cleavage are 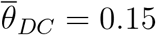 rad and *s*_*MW*_ = 1.27. (**H**) Example of active re-orientation of a nucleus immediately prior to cell division. (**I**) Schematic of division induced defect climb along parasegments.

Once a PSPR is assembled, its constituent cells characteristically undergo two rounds of highly choreographed cell divisions. Within each parasegment, mitotic waves are initiated at the ventral midline and spread outward towards the dorsal regions of the embryo (Fig. 2B). The timing of these intra-segment waves is stereotypic among all segments (Fig. 2D) and, since PSPRs are constructed sequentially, their onset is staggered between adjacent PSPRs (Fig. 2, E and F). This choreography leads to two orthogonal phase waves of cell divisions with distinct wave velocities: a fast wave within each PSPR spreading ventral-to-dorsal and a slower one across PSPRs that moves anterior-to-posterior (Fig. 2C). After completing these two mitotic waves, each PSPR subsequently undergoes rapid differential cleavage in localized regions adjacent to the ventral midline (Fig. 2B). Together, these results suggest that PSPRs behave as weakly coupled, independent units running the same modular proliferation program. In other words, cells in different PSPRs begin to divide at different times, but the relative timing of division waves is shared among segments.

The orientations of the divisions comprising the mitotic waves are tightly distributed about the A-P axis (Fig. 2G). This coherence of cell division axes appears to be actively maintained. We frequently found that condensed nuclei with the wrong orientation would rapidly rotate to align with the global division axis (Fig. 2H). This patterning of oriented and wave like timed cell divisions ensures that the mitotic waves manifest a kind of ‘defect climb’ (Fig. 2I). The mitotic waves gradually insert new rows into the bulk of the grid. Incomplete rows manifest defects at their left and right edges (Fig. 1N). As the mitotic waves unfold, these defects are pushed out towards the dorsal regions of the tissue, leaving behind an intact fourfold ordered grid. Defect climb in non-living systems typically only becomes the dominant mode of defect motion at extremely high temperatures [28]. Here, the presence of cell divisions excites defect climb at room temperature, such that defects, which would otherwise disorder the lattice, are healed by subsequent divisions. In this way, cell divisions are able to serve as an order generating mechanism.

Next, we investigated how the division choreography dynamically shapes the ectoderm at the tissue scale. For the purposes of this analysis, we focused on a subset of six PSPRs. We determined that growth proceeds in two stages (Fig. 3A). In the first stage, mitotic waves extend the germband by inserting new rows without changing the average cell density (Fig. 3, B and C). The tissue elongates along the A-P axis and increases in total area, but its width remains approximately constant. In the second stage, the tissue undergoes convergent extension. Its width is sharply pinched, but its length continues to increase in such a way that the total tissue area is held approximately fixed. Since the rate of cell division remains constant throughout both stages (Fig. 1F), the average cell density necessarily increases during convergent extension.

**Fig. 3.**
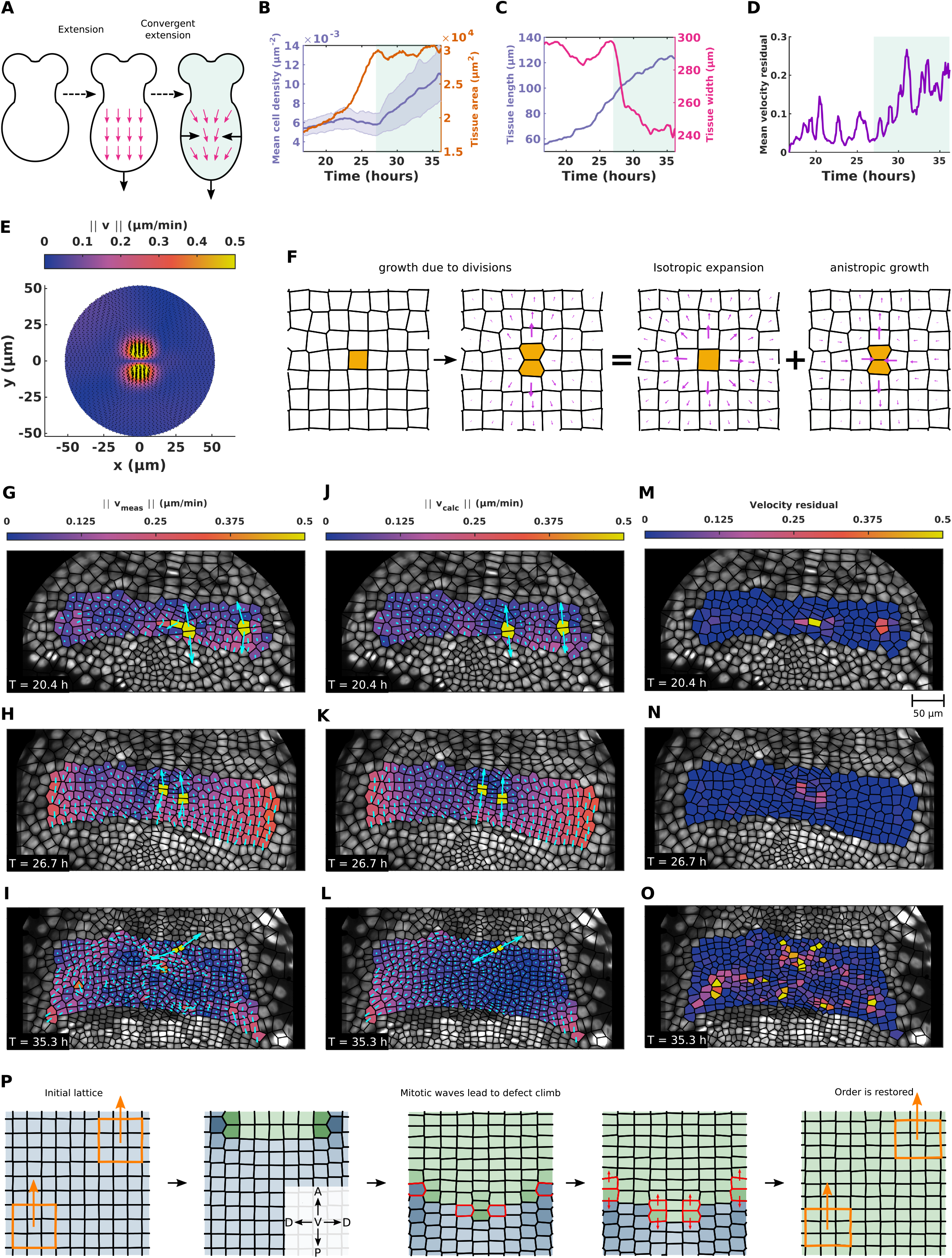
Active cell divisions dictate tissue velocities during germband extension. (**A**) Schematic of the two stages of tissue scale growth observed in the germband. (**B** and **C**) Tissue scale observable fields in the germband. Tissue cartography ensures 3D geometry is properly accounted for. Shading corresponds to the two observed stages of growth. (**B**) Mean cell density and total tissue area. Blue shaded region shows standard deviation. (**C**) The tissue length measured along the A-P axis and the tissue width measured along the D-V axis. (**D**) Mean residual between the measured cell velocities and the cell velocities predicted by the active hydrodynamic model. (**E**) The magnitude of the velocity of the mean cell division event. Arrows showing orientation are scaled by the norm of the velocity. Schematic of the flow field induced by a single division event. (**G** to **I**) Cell velocities measured by single cell tracking. (**J** to **L**) Predicted cell velocities. (**M** to **O**) Single cell velocity residuals for the measured and calculated velocity fields shown in (G to I) and (J to L), respectively. (**P**) A schematic summarizing the role of division choreography in generating and maintaining orientational order. Hue corresponds to number of cell divisions within a lineage and saturation and value represent orientational order (less ordered regions are darker). Order is maintained and the grid is restored so long as the divisions are oriented along a single axis and each cell divides once before any particular cell divides twice. Cells divisions shown in the middle panels are highlighted in red. Orange regions in the first and last panels show the direction of local order is coherent over long distances and preserved by the division choreography.

These two stages also feature markedly different dynamics of the global order parameter (Fig. 1J). During the first stage, when the large scale motion of the tissue appears to be dominated by the mitotic waves, the global order parameter consistently increases. In the second stage, wherein the tissue experiences a reduction in width that is not explained by the mitotic waves alone, the global order parameter is initially static, but slowly falls off.

Having separately elucidated the kinetics of cell proliferation and the time course of tissue geometry, our next goal was to explicitly connect local cell behaviors with global tissue shape and order. We adopted a coarse-grained description of the tissue-scale mechanics, similar to recent applications aimed at decoding how active forces generate tissue flow patterns [29–33]. Motivated by our observation of exponential growth through cell proliferation, we developed a hydrodynamic model, similar to the Stokes equations, that directly links tissue flow to bulk contributions from oriented cell divisions via an active sourcing term (see supplementary text). During division, the mitotic spindle within cells generates extensile forces akin to a force dipole [13, 34]. These division events push on nearby cells, deforming the dividing cell’s neighbors and generating local plastic strain. Viscoelastic relaxation of this instantaneous strain leads to flow in the surrounding tissue (Fig. 3F). In this way, many division events can combine to create large scale collective motion. We investigated the extent to which the integrated deformations induced by divisions could collectively account for the observed global tissue flow. We first deduced the average plastic deformation induced by a single cell division by rotating all observed cell division events into a common frame (Fig. 3E, fig. S12 and fig. S13). Informed by the features of this average flow field (fig. S12 and fig. S13), individual division events were modeled as circular inclusions, i.e. finite regions within which the tissue undergoes a permanent plastic strain [35], with an orientation chosen to align with each measured division axis. We then solved our model using the finite element method (FEM) with Dirichlet boundary conditions to predict the tissue flow from observed cell divisions.

To benchmark accuracy of this model, we compared the predicted tissue flow to velocity fields quantified from individual cell tracking. Measured tissue flow fields were mostly harmonic, irrotational, and divergence-free, except at cell divisions, where the divergence, curl, and Laplacian of the flow fields all spiked (Fig. 3, G to I and fig. S11). Visually, the flow fields predicted from our model appear strikingly similar (Fig. 3, J to L). The velocity residual quantitatively confirms that our simple model, with only one free parameter, accurately predicts both the direction and magnitude of the observed tissue flow (Fig. 3, M to O) (see supplementary text). During the first phase of growth, velocity residuals are typically below 10% (Fig. 3D). This suggests that about 90% of the flow can be accounted for in terms of cell divisions as the dominant bulk contribution. Velocity residuals subsequently increase moderately during convergent extension, reaching a typical level around 20%. This suggests divisions are still the main driver, but other mechanisms, currently not accounted for by the model, provide small but measurable adjustments.

In this work, we combined mathematical modeling with quantitative flow analysis to show that cell divisions are the primary drivers of global tissue flow during *Parhyale* germband extension. We uncovered a global fourfold bond orientationally ordered phase that emerges via precisely oriented cell divisions. In a scheme we call defect driven morphogenesis, cell proliferation introduces defects into the local tissue structure, which are then healed by subsequent cell divisions, ultimately giving rise to a highly ordered cell network (Fig. 3P). The choreography of these events is arranged in a timed mitotic wave, spreading at distinct wave velocities across the A-P and D-V axes. The anisotropic timing of this wave of cell divisions results in defect climb at room temperature. Defects migrate out of the ectodermal bulk toward the boundary, leaving behind an intact lattice. Defect driven morphogenesis is both an efficient and a highly robust mechanism for establishing global orientational order in presence of cell divisions. Similarly to non-living matter [36], the insertion of new particles is an efficient strategy for exploring regions of configuration space corresponding to orientational order. Provided a preferred axis, order can then be produced by having cells divide according to independent, internal timing mechanisms. This timing does not have to be precise (fig. S5). Order within a local region is preserved so long as most of the cells within that region divide once before any particular cell divides twice (Fig. 3P).

We showed that defect driven morphogenesis at the single cell level relies on tightly oriented cell divisions. Our analysis demonstrated that neither the orientation (fig. S7) nor the timing of cell divisions (fig. S8 to S10) exhibit strong correlations with mechanical or geometric signals. This suggests the existence of biochemical signals, such as morphogen gradients or planar cell polarity, that actively instruct the cell division axis orientations. For small direct developers, like *Parhyale*, arranging cells in an ordered grid might be one of very few possibilities to establish a coordinate system in the presence of growth [19]. It is intriguing to speculate whether large embryos, with abundant cell numbers, utilize a similar ordering strategy when arranging cells in bigger periodic units, such as the somites in vertebrates [37]. Future work will investigate how defect driven morphogenesis differs in implementation between relatively small embryos and embryos with large cell numbers.

## Acknowledgments

S.J.S. acknowledges NIH grant 5R35GM138203-02. This work was initiated during the 2016 Santa Barbara Advanced School Of Quantitative Biology, supported by NSF Grant No. PHY-1748958, NIH Grant No. R25GM067110, and the Gordon and Betty Moore Foundation Grant No. 2919.02. M.J.B. acknowledges support from the NSF through the Materials Science and Engineering Center at UC Santa Barbara, DMR-1720256 (iSuperSeed). We thank A. Pavlopoulos, C. Wolff, B. Shraiman, and Streichan lab members for fruitful discussions. We thank A. Pavlopoulos for sharing transgenic *Parhyale* culture and methods used to generate the data recorded for this manuscript.

## Materials and Methods

### Light sheet microscopy

For live imaging of transgenic parhyale embryos, we utilized a custom built MuVi SPIM [1]. This microscope has two excitation and two detection branches. Both used water dipping objectives (App LWD 5x, NA 1.1, Nikon Instruments Inc. for detection, and CFI Plan Fluor 10x, NA 0.3 for excitation). Furthermore, each detection branch consisted of a filter wheel (HS-1032, Finger Lakes Instrumentation LLC), with emission filters (BLP02-561R25, Semrock Inc.), tube lens (200 mm, Nikon Instruments Inc.) and a camera (sCMOS - Hamamtsu Flash 4.0 V2), with effective pixel size of 0.262 mm. The illumination branches featured a tube lens (200 mm, Nikon Instruments Inc.), scan lens (S4LFT0061/065, Sill optics GmbH and Co. KG), galvanometric mirror (6215 hr, Cambridge Technology Inc.), and discrete laser line (561LS OBIS 561nm). Optical section employed a translation stage from Physik Instrumente GmbH and Co. KG (P-629.1CD with E-753 controller), a rotation stage (U-628.03 with C-867 controller), and a linear actuator (M-231.17 with C-863 controller).

### Data post processing and microscope automation

To operate the microscope, we used Micro Manager [2], installed on a Super Micro 7047GR-TF Server, with 12 Core Intel Xeon 2.5 GHz, 64 GB PC3 RAM, and hardware Raid 0 with 7 2.0 TB SATA hard drives. For each sample we recorded 4 views, separated by 90*°* rotated views, with optical sectioning of 2 *µ*m, and temporal resolution of 5 min. We embedded the embryos in agarose containing beads as a diagnostic specimen. This was used to register individual views into a common frame by utilizing the Fiji multi view deconvolution plugin [3], resulting in a final image with isotropic resolution of .2619 *µ*m.

### Extraction of dynamical surfaces of interest

The output of the lightsheet microscope is a time series of 3D grids whose voxel values correspond to intensity of the nuclear label. Extraction of the dynamical surface of interest from these data sets was performed in two stages: (1) 3D surface extraction and (2) 2D pullback map construction. In the surface extraction stage, the volumetric data of a representative time point was classified over the nuclear label using the machine learning software Ilastik [4]. The resultant probability map was then fed into MATLAB and a static surface of interest was extracted using the morphological active contours method [5] (fig. S15), a type of level-set based segmentation algorithm well suited to segmented complicated, closed surfaces. The output of this segmentation is a 3D binary level-set, with identical dimensions to the data, where ‘1’ values corresponded to the interior of the closed surface (all embryonic tissue and yolk) and ‘0’ values corresponded to regions external to the *Parhyale* egg. The boundary of this binary level-set is point cloud, a subset of which included voxels corresponding to the embryonic tissue. This point cloud was subsequently triangulated using Poisson surface reconstruction [6]. The result was a topologically spherical mesh triangulation.

In the next processing step, this static surface was used as a seed to extract the dynamically changing surface at each time point. Recall that at this developmental stage the embryonic tissue is a topological disk sitting on top of a spherical yolk. The embryonic tissue was therefore contained in a disk-like subregion of the sphere-like surface triangulation. In order to extract this region of interest, the entire sphere-like mesh was mapped into the plane using the orbifold Tutte embedding method [7]. This method generates a topologically consistent parameterization of the sphere in the plane allowing us to view the entire surface at once with minimal geometric distortion. Next, a static submesh of the region of interest on the static surface was selected by hand using the orbifold pullbacks. Although static, this region of interest was large enough that it contained all relevant sections of the embryo as it grew and deformed over time. A set of ‘onion layers’ was then created by displacing the submesh along its positive and negative normal directions. A stack of pull-back images was then created for each time point with one image in the stacks for each displaced onion layer. The number of layers and the inter-layer spacing was chosen so that all of the geometric features of the dynamic surfaces were captured for the various time points somewhere within the image stack. These stacks were then fed back into Ilastik and batch processed again over the nuclear label. The result was a time dependent field of normal displacements over the static seed surface that transformed the static surface into the corresponding dynamic surface for each time point. These dynamic triangulations of the evolving region of interest were then separately mapped into the unit disk conformally via Ricci flow [8]. Such a conformal mapping is only unique up to a Möbius automorphism of the unit disk. In other words, unless care is taken to register the pullbacks, the resultant images may be wildly misaligned in pullback space from time point to time point. With this in mind, the time series of conformal pullbacks was iteratively registered to fix the conformal degrees of freedom within the pullbacks. Essentially, corresponding mesh vertices at subsequent times were approximately matched in 2D by finding an optimal Möbius automorphism of the unit disk that registered as many points as possible without sacrificing the conformality of the parameterization [9]. The final result was a sequence of maximally aligned conformal pullbacks of the growing embryo to the plane.

It should be noted that tissue cartography framework allows the user to implement not only conformal mappings, but a broad variety of parameterizations depending on the application. The visualization of the full embryo dynamics in Movie S3 was created using an ‘As-Rigid-As-Possible’ parameterization [10]. This parameterization attempts to produce a high quality visual representation of the 3D surface in the plane by settling on an optimal balance of angle distortion and area distortion.

### In-plane cell segmentation and pathline reconstruction

One primary benefit of tissue cartography is that processing data in low dimensions greatly reduces the computational complexity of various analysis procedures. We exploited this benefit by segmenting nuclei directly in the 2D pullback images. Images were first classified in Ilastik. The resultant probability maps were then fed in MATLAB where the nuclei were segmented using a custom built version of the watershed algorithm [11]. Custom additions to MATLAB’s built-in `watershed` functionality were necessary to account for spatially proximal nuclei which were initially undersegmented, i.e. many nuclei were counted only as a single object. Special care was taken during this step in adjusting watershed parameters to ensure that adjacent nuclei were properly distinguished from each other.

Once segmented, nuclei were tracked semi-automatically using an enhanced point-matching procedure. For a given time point, the input to this procedure included a pullback image and segmentation at time *t* and another subsequent pullback image and segmentation at time *t* + 1. First, the subsequent pullback image was registered onto the previous image using the Demon’s deformable image registration algorithm [12]. The resultant displacement fields were then applied to the nuclei locations at time *t* + 1. Point matching was then used to associate the nuclei locations at time *t* with the displaced nuclei at *t* + 1. Displacing the nuclei locations at *t* + 1 to align more closely with the locations at *t* reduced discrepancies in the point matching associated with large nuclear motions. This process was applied iteratively until pathlines were generated for all cell lineages at all time points. Despite the enhancements, some manual correction was still necessary. Manual corrections were applied using a custom built MATLAB GUI. The pathlines were outputted as a digraph where nodes represented particular cells at particular time points and edges stored the information about the temporal relationships between nodes. Cell divisions could then be extracted from this tracking structure by locating events where single tracks split into two lineages.

Another benefit of tissue cartography is that the geometric information reflecting the fact that in-plane dynamics are occurring on a 2D surface embedded in 3D space are properly preserved. In particular, knowing the 2D locations of nuclei in pullback space provides an explicit correspondence to their locations on the surface in 3D. Therefore, once the tracks were constructed in 2D, it was trivial to extract full 3D nuclear pathlines. Velocities were constructed as simple backward differences between 3D nuclei locations. A backward difference was used since forward differences can generate ambiguities at cell division events where the forward difference velocities of the children sum to zero. 3D velocities were decomposed into tangential and normal components relative to the dynamic surface. Tangential velocities were then consistently transformed back into pullback space for display purposes using a discretization of the Jacobian on mesh triangulation faces.

### Calculation of discrete curvatures

The discrete curvature was calculated for each dynamic mesh triangulation as a function of time using standard discrete constructions (fig. S14) [13]. The Gaussian curvature *K*(*υ*_*i*_) of a mesh vertex *υ*_*i*_ was taken to be

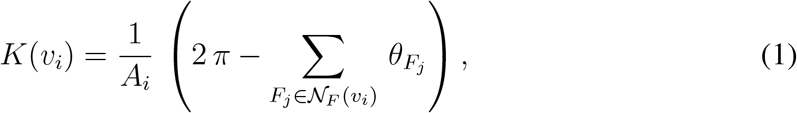

where *A*_*i*_ is the area associated to each vertex via barycentric subdivision of the triangles attached to that vertex, 𝒩_*F*_ (*υ*_*i*_) is the set of incident faces *F*_*j*_ attached to *υ*_*i*_, and 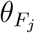 is the internal angle of face *F*_*j*_ corresponding to *υ*_*i*_. The mean curvature *H*(*υ*_*i*_) of a vertex *υ*_*i*_ was calculated according to

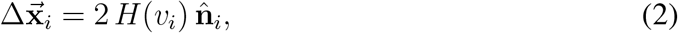

where 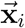 is the 3D location of vertex *υ*_*i*_, 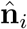 is the unit normal vector corresponding to vertex *υ*_*i*_, and Δ denotes the Laplace-Beltrami operator. The discrete Laplace-Beltrami operator was implemented using the familiar cotangent discretization [13]

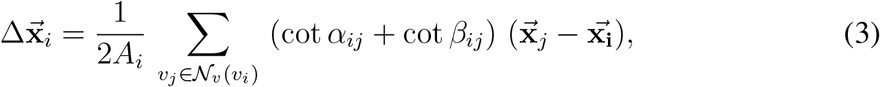

where 𝒩_*υ*_(*υ*_*i*_) is the neighborhood of vertices attached to vertex *υ*_*i*_ and *α*_*ij*_ and *β*_*ij*_ are the two internal angles of the triangles opposite the edge shared by vertex *υ*_*i*_ and *υ*_*j*_. In order to extract *H*(*υ*_*i*_) from Eq. (2), it was first necessary to calculate 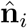, which was taken to be the angle-weighted average of the face unit normal vectors of the triangles incident to vertex *υ*_*i*_ assuming a counter-clockwise orientation of vertices in the faces. This choice breaks the degeneracy in the orientation of the unit normal and allowed for simple extraction of a signed mean curvature. The panels in fig. S14, were constructed by averaging together the Gaussian and mean curvatures, respectively, of all vertices found to lie within a particular Voronoi polygon corresponding to a specific cell after mapping the dynamic meshes conformally into the plane.

### Construction of correlation functions

In order to construct the two-point orientational order correlation functions, we first calculated the fourfold and sixfold orientational order parameters for each cell at a particular time point (Fig. 1F and fig. S1). Next, for each pair of cells, denoted here by their locations **x**_1_ and **x**_2_, we calculated the intercellular distance. This distance was taken to be the geodesic distance along the dynamic surface between the locations of the cell centroids on the 3D mesh triangulation [14]. We also calculated the product 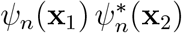 for each pair of cells, where *n* = {4, 6}. We then partitioned the intercellular distances into a set of bins. All pairs whose spacing lay between *r* and *r* + *dr*, where *dr* was the width of a bin, were then averaged together to calculate the two-point orientational order correlation function 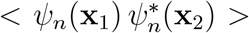. It was assumed for this calculation that this quantity depended only on the scalar distance *r* between pairs of cells, i.e. 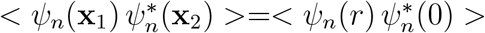. Note that under averaging, only the real part of the product 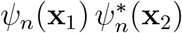 contributed, since 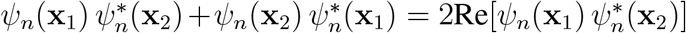. Finally, the intercellular distances distances were normalized by an average cell length scale, calculated as the square root of the average area contained by each cells Voronoi polygon mapped back into 3D.

Construction of the pairwise radial distribution function *g*(*r*) proceeded similarly and also relied on the calculation of the pairwise geodesic distance between cells. Following previous work [15, 16], we constructed multiple distribution functions along cuts with specific orientations, rather than simply considering all pairs of cells with spacing between *r* and *r* +*dr*. In particular, we considered cuts along the A-P and D-V axes. All pairs whose relative orientation lay close to these cuts were consolidated into histograms within each bin segmenting the intercellular spacing. In this way, we managed to capture the anisotropy in the distributions. The final quantity reported as a function of *r* is the number of cell pairs in a particular bin along the cut divided by the total number of cells present at that particular time.

### Circular statistics for division events

In order to properly analyze the distributions of division events, it was necessary to construct measures of statistical properties that properly accounted for the nematic nature of division events, i.e. a division event with orientation *θ* is physically identical to a division event with orientation *θ* ± *π*. Extending familiar measures of circular distributions [17], the modified circular mean of the orientations of a set of division events, {*θ*_*n*_} where *n* = 1, … *N*, was defined to be

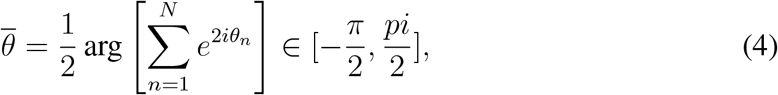

which is invariant under the transformation *θ*_*n*_ →*θ*_*n*_ ± *pi* for any *θ*_*n*_. If the division orientations are tightly distributed around a single value, then this quantity will also be close to that particular value. We also define a modified measure of angular dispersion

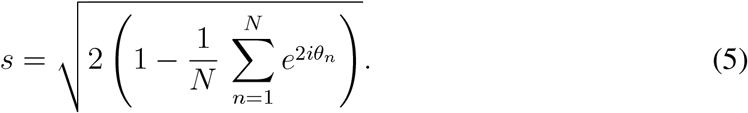

Note that this is a dimensionless quantity. Our measure of angular dispersion varies between *s* = 0 for perfectly oriented divisions (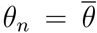 for all *n* and 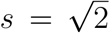 for totally isotropic divisions.

### Determination of the average division velocity

Division events were extracted from the tracking structure and the resulting sample set was pruned for quality (sufficiently far from the boundary of the tissue, sufficiently far from another division event, etc). In order to compare different events, divisions were translated and rotated so that the center of mass of the daughter cells lay at the origin and the division axis lay along the *y*-axis. The measured velocities of the daughter cells and of cells in the third-order natural neighborhood of the daughter cells were then interpolated onto a fine mesh triangulation using generalized Hessian-energy scheme that minimizes distortion in the interpolated field at the boundary of the triangulation [18]. The interpolated velocity fields were then averaged across division events to find the mean velocity induced by divisions (fig. S12). Gradients of the resultant velocity fields were calculated on the mesh triangulation using a custom built implementation of the Discrete Exterior Calculus (DEC) [19] in MATLAB (fig. S13).

### Numerical prediction of tissue velocities from cell divisions

An active hydrodynamic model was used to predict the tissue velocities resulting from the collective motion induced by cell divisions. The full details of the model are presented in the supplementary text. The model was solved numerically for actual data using a custom built machinery based on the DEC. First, a fine mesh triangulation was constructed over a subset of tracked cells at a particular time (fig. S16A). Circular holes containing cells about to divide were then removed from the triangulation. These holes represented the finite-size circular Eshelby inclusions used to model the velocities induced by cell divisions (supplementary text). The size of the holes was chosen so that the area of the circular holes equaled the area of the Voronoi polygons of the corresponding cells. Removing the inclusions from the mesh renders the task of predicting velocities a simple boundary value problem with Dirichlet boundary conditions. The velocities on the interior boundary vertices were set to match the analytical predictions of the model. The velocities on the exterior boundary were set to the measured velocities. Using the measured velocities on the exterior boundary captured how cells not included in the subregion contributed to the relevant motion within the region over which the velocities were predicted. The results of the numerical solution was a velocity vector for each triangulation vertex. For cells that did not divide, all vertices contained within each corresponding Voronoi polygon were then averaged to produce a single cell velocity vector. Note that many vertices were averaged for each polygon, so that prescribing Dirichlet boundary conditions on the triangulation did not correspond to prescribing cell-scale velocities. Note that under deformation the circular inclusions were deformed into ellipses. The locations of the foci of these ellipses were given directly by the fluid mechanical model. The velocity of cells that divided were set to be the displacement of these foci from the center of the undeformed circular inclusion.

### The velocity residual

To compare the measured flow fields **v**(**x**) to the predicted flow fields **u**(**x**) in a quantitative fashion, we defined a global measure for the spatial velocity residual that was insensitive noise dominated fluctuations in the regions of slow flow. Let

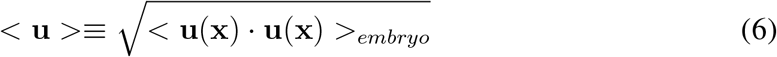

define an overall magnitude of the field **u**(**x**). Here, < **u**(**x**) ·**u**(**x**) >_*embryo*_ denotes an average of the spatially dependent field **u**^2^(**x**) = **u**(**x**) *·***u**(**x**) over the entire embryo and is therefore not space dependent. We define our velocity residual as

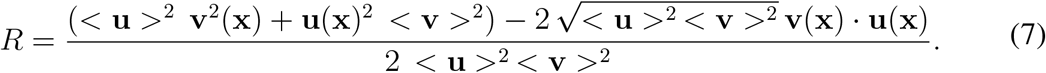

This residual provides a spatial discrepancy map, indicated the prediction quality as a function of location on the embryo. An identical velocity residual was used in [20].

### Generation of synthetic data sets and comparison with measured data

The synthetic data sets were generated to understand the extent to which the measured order was statistically significant and how the finite size of the germband effected the order. First, the length and width of the germband at each time was extracted. These were determined by tagging representative rows and columns of cells, following these rows and columns over time, and calculating the geodesic length along each row and columns. The average geodesic length of the rows (columns) was taken to be the width (length) of the germband at a particular time. We also extracted a mean cell density at each time point by averaging the inverse 3D areas of cells. One thousand synthetic data sets were generated for each time point within rectangles with the same length and width as the germband. Points representing cell centroids were generated to exhibit the same cell density as the germband using a fast Poisson-disk sampling method [21]. The connectivity of these randomly generated points was approximated using Voronoi tesselation. This connectivity allowed for the calculation of the order parameters and orientational correlation functions over the finite size samples. We also calculated the radial distribution functions along the A-P axis (length of the rectangle) and the D-V axis (width of the rectangle) for each synthetic data set (fig. S3) using the same method of construction as for the measured data.

Additionally, we compared the corresponding measured and synthetic distributions of orientational order parameters according to the two-sample Kolmogorov-Smirnov (K-S) test [22] using MATLAB’s `kstest2`. This implementation of the K-S test returns two measurements that assess the confidence with which one can assert that two sets of observed random variables are drawn from the same distribution. The first is the K-S statistic, which is simply the maximum difference between the empirical CDFs of the two sample sets. The larger the K-S statistic, the greater the discrepancy between the two sample sets. The second measurement is the asymptotic *p*-value, which is the probability of observing a test statistic as extreme as, or more extreme than, the observed value under the null hypothesis that both samples are drawn from the same distribution. All K-S statistics and *p*-values are reported in Table S1. At the confidence level *α* = 0.05, the null hypothesis is rejected for all distributions, indicating it is unlikely that the observed order at any time point is due to chance. We note that the *p*-values for fourfold order at intermediate and late times are vastly smaller than the corresponding *p*-values at early times, while the *p*-values for sixfold order do not change as drastically. This implies that while it is still improbable that the observed fourfold order at early times is due to chance, it is hugely more likely when compared to intermediate and late times after the order generating choreography has unfolded.

Similar methods were used to generate the synthetic illustration of a translationally ordered system (fig. S4). We patterned perfect square lattices on rectangles with the same length and width as the germband for the same representative time points. The lattice spacing was set to match the measured cell density at the corresponding time point. We then generated one hundred synthetic data sets for each time point by adding Gaussian white noise to the lattice site positions with a signal-to-noise ratio of 5. Calculation of the orientational order parameters, the orientational correlation functions, and the radial distribution functions were performed in the same manner as the other synthetic data sets.

## Supplementary Text

### Fluid mechanical model

We investigated the role of cell divisions in producing tissue scale flow fields through the use of an active hydrodynamic model. Continuum mechanics is an effective method for capturing how force balance is associated with local deformation and flow [23]. In the simplest models, the forcing that generates flow can be neatly decomposed into a set of separate passive and active contributions. The tendency of cells to resist deformation and the viscous coupling between cells translate local forces into long range flows. Interactions among these various contributions, both locally and over large scales, can be then be built up with increasing degrees of complexity to create more accurate descriptions of biological realities. This framework has proved effective in numerous biological applications elucidating how active forces generate non-trivial flow fields [20, 24, 25].

The tissue scale mechanics of epithelia is best understood as emerging from the collective mechanics of the tissue’s constituent cells [26]. The cell scale mechanics are dominated by the influence of intracellular cytoskeletal cortices [27, 28]. These structures are capable of supporting stresses throughout the cell’s bulk interior and can deform the cell either actively or in response to external stresses. The individual cortices of neigboring cells are coupled together by cadherin mediated adherens junctions [29] into a global trans-cellular mechanical network. Crucially, these networks are both ‘active’ and ‘adaptive’. Active contributions, such as cell divisions [30] or the contraction of trans-cellular actomyosin networks [20], act both locally and over large scales to deform the tissue. These active deformations generate passive stress distally from the activity via the coupling between cells. Stresses created in this fashion can then be relaxed either locally, via internal rearrangement of the cytoskeletal cortex [31, 32], or at the tissue scale, via cell rearrangements [24]. The result is an exotic type of viscoelasticity, wherein the tissue responds elastically to active stresses over short timescales and then reverts to a fluidlike flow at longer timescales as the tissue adaptively relaxes stress. When the timescale of growth (i.e. rate of cell division, etc.) is long compared to the timescale of mechanical relaxation, the tissue essentially behaves like a slowly creeping fluid in quasistatic mechanical equilibrium with both the active and external forces. For our purposes, a detailed description of the complex cellular processes mediating the active stresses and adaptive relaxation is unimportant. Instead, we adopt an approximate continuum scale description of the tissue that captures the relevant behaviors of the system.

At short time scales, the tissue behaves elastically [33, 34]. Incremental increases in strain generate corresponding increases in stress. These stresses are then subsequently relaxed over a time scale *τ*_*R*_. The total stress ***σ*** = ***σ***^*e*^ + ***σ***^*a*^ can be decomposed into a sum of the elastic stresses ***σ***^*e*^ and the active stresses ***σ***^*a*^. For simplicity, we assume that the tissue is a flat two-dimensional surface. Supposing the tissue behaves like a Maxwell viscoelastic fluid, the time-evolution of the elastic stress is given by

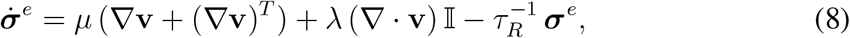

Where 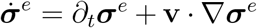 denotes a convective time derivative, **v** is the local velocity or rate of displacement, and *µ* and *λ* are the Lamé parameters characterizing a linear isotropic stress-strain relationship. The Lamé parameters are assumed to be spatially homogeneous, but may depend on time. A more complete description would include the corotational time derivatives of ***σ***^*e*^ on the left hand side, but we omit them for simplicity. The first two terms on the right hand side describe the generation of stress in proportion to the rate of strain. The final term parameterizes the relaxation of the elastic stress. We have made the assumption that 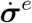 only depends on ***σ***^*a*^ implicitly through the final term 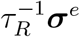 (see the force balance condition below). This is not meant to imply that adaptive cell behaviors do not actually depend explicitly on the active stresses. It is merely a mathematical simplification. In reality, there may exist a complex interplay between mechanics, gene expression/cell fate, and adaptive cellular behaviors, but this lies beyond the scope of this work.

In general, the tissue flow velocity can be described by the Cauchy momentum equation [23]

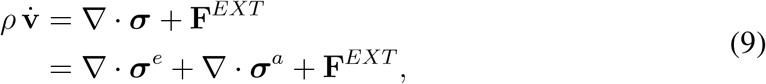

where *ρ* denotes the material density and **F**^*EXT*^ denotes any external body forces acting on the system. If the rate of mechanical relaxation is sufficiently fast to reach quasi-equilibrium, 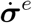 vanishes, leaving us with

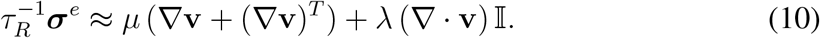

Substituting this into Eq. (9), we find that

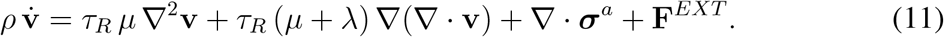

In other words, the transient elasticity of the system gives rise to fluidlike behavior in the quasi-stationary regime parameterized by a set of effective viscosities. The effective shear viscosity *ν*_1_ ≡ *τ*_*R*_ *µ* tends to resist shearing motion, while the effective bulk viscosity *ν*_2_ *≡τ*_*R*_ (*µ*+*λ*) resists the isotropic compressible part of the flow. If the effective viscosities are sufficiently large and the corresponding motion is sufficiently slow we can neglect the inertial term relative to the viscous terms 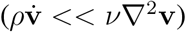. If, for simplicity, we furthermore assume that there are no external forces acting on the system, we are left with the following force balance condition defining the tissue flow velocities in terms of the active stresses

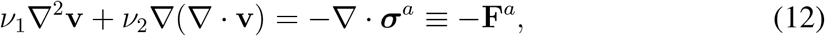

which can be solved given suitable boundary conditions.

### Cell divisions and active stresses

In general, there may be numerous different types of active stresses acting within the tissue during growth [20, 30, 35]. These various sources of active stress will likely not contribute with equal importance to the shaping of the nascent tissue. Our observations of the cell division choreography indicate that cell proliferation is the primary factor mediating tissue velocities. In order to simplify our model, we assume that the only relevant active stresses are due to these cell divisions.

Our first consideration in modeling active stresses due to cell divisions is to understand how division events might be incorporated into a spatiotempoally coarse grained scheme. With respect to timing, daughter cell separation in real cell division events occurs over a short, but finite time [36]. We ignore the complexity of the rapid subcellular properties mediating mitosis and model cell divisions as instantaneous events (for our purposes any process that occurs faster than 5 min, the time resolution of our microscope data, is considered instantaneous). Real cell divisions push on the surrounding tissue, which responds elastically over the short time scale of the actual division, before remodeling to relax the resulting stress. In our model, both the plastic strain due to a division and the subsequent relaxation are assumed to occur instantaneously so that the fluid is always in quasistatic mechanical equilibrium. With respect to spatial coarse graining, we note that the length scale of a single cell division is small compared to the scale of the entire germband. Numerical experimentation showed that approximating division events as point force dipoles produced inaccurate flow fields. We therefore model divisions as small, but finite size events to regularize this inconsistency. We also make the approximation that the inclusion lives in an infinite, otherwise quiescent medium in order to make analytic progress.

The mathematical structure of the fluid mechanical equations of motion in Eq. (12) is identical to the structure of the Navier equations of linear elasticity. Exploiting this similarity, we can model the instantaneous displacement (read velocity) due to a cell division as that of an Eshelby inclusion. In the context of classical elasticity, an Eshelby inclusion is a finite subregion of an elastic body which undergoes a permanent plastic deformation [37]. Eshelby inclusions, and the associated theories of elastic multipoles [38], have proved useful in numerous applications including understanding shear localization in amporphous solids [39] and the properties of mechanical metamaterials [40]. In the current context of viscoelastic tissue growth, this displacement corresponds to both the plastic deformation and the subsequent viscous relaxation, which are assumed to both occur instantaneously on time scales relevant to growth. Relying on the well-trodden mathematical history of Eshelby inclusions, we may, in the following derivation, use terminology similar to Eshelby’s original exposition for elastic materials (e.g. eigenstrain). For clarity, we emphasize again that we are modeling the tissue as a viscoelastic fluid, not an elastic solid, and that the parameters of our model are effective viscosities, not elastic moduli which would vanish in an orientationally ordered phase.

In particular, we choose to model the division as a circular Eshelby inclusion of radius *a* and eigenstrain 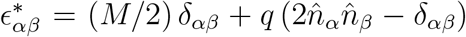, where *δ*_*αβ*_ is the Kronecker delta and 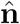 is a unit vector along the division axis. Note that we have adopted an index notation to better account for the high-rank tensorial nature of the following calculations. Greek indices vary in the set {*x, y*}. We also adopt the Einstein summation notation so that all repeated indices are summed. It was shown by Eshelby that the constraining medium generates a constant strain within the inclusion 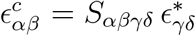, where *S*_*αβγδ*_ is a constant tensor for any elliptical inclusion. For a circular inclusion

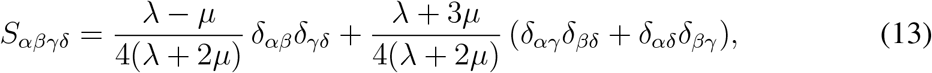

which yields the following constrained strain within the inclusion

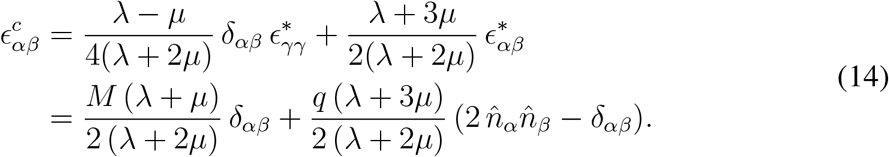

Thanks to the linearity of the system, the corresponding velocity, i.e. instantaneous displacement, within the inclusion can now be computed as

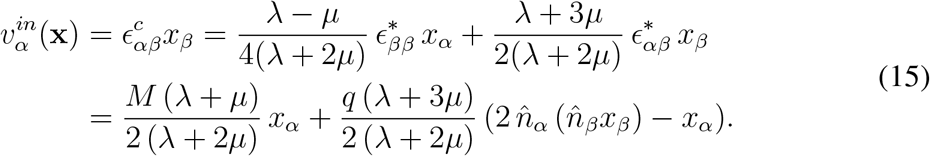

The velocity outside of the inclusion will satisfy the biharmonic equation 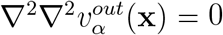, subject to continuity at the surface of the inclusion and must also tend to zero as *r* → ∞ where *r* ≡ || x ||. Recalling the radial solutions of the biharmonic equation in 2D (i.e., 1, ln *r, r*^2^, and *r*^2^ ln *r*), we construct the general solution by considering all combinations of derivatives of the radial solutions that are linear in the eigenstrain, tend to zero at infinity, and transform like a vector field:

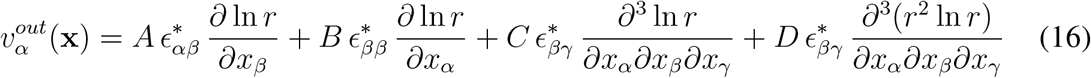

where *A, B, C*, and *D* are constants to be determined. Recall that the velocity outside of the inclusion is a solution to the equation

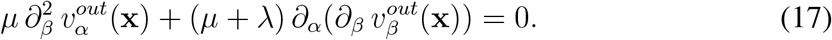

The fact that 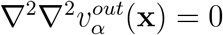 is a necessary, but insufficient condition on any solution of Eq. (17). With this fact in mind, we begin to calculate the constants by first computing

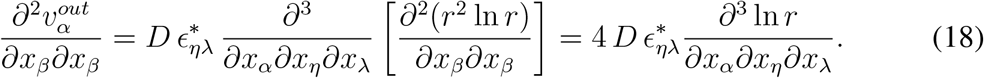

In the first equality, we used the identity

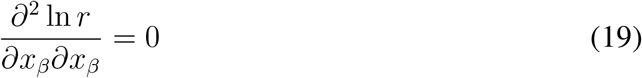

to cancel the terms proportional to *A, B*, and *C*. In the second equality, we made use of the fact that

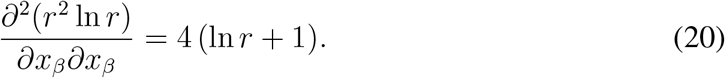

Next we compute

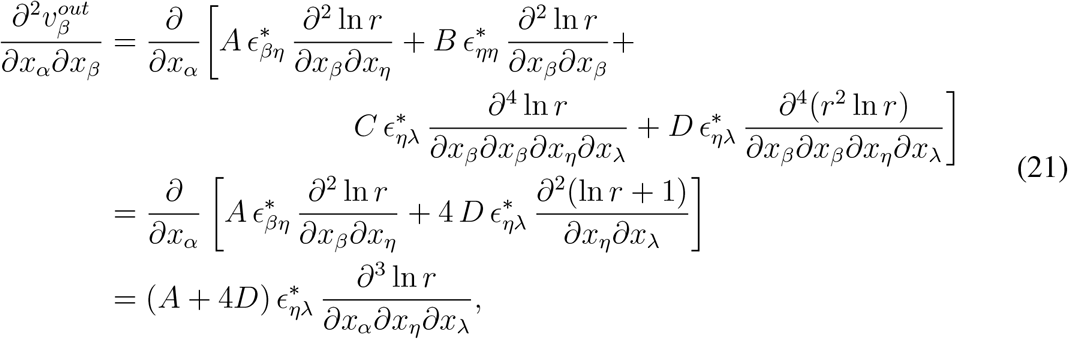

where we have once again made use of both Eq. (19) and Eq. (20). Substituting these results into Eq. (17) yields

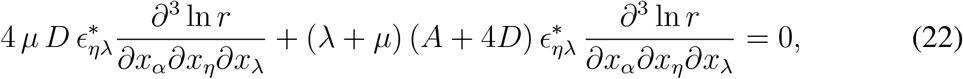

which, when simplified, reveals that

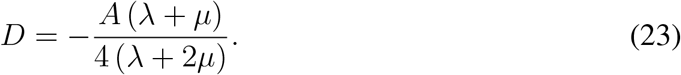

The velocity outside the inclusion can therefore be written as

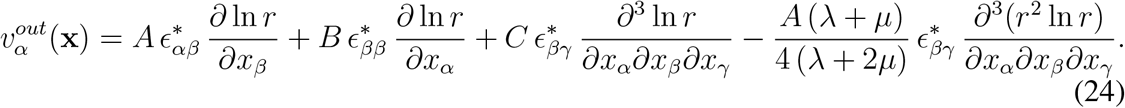

The following identities are now required to make further progress:

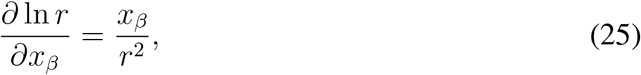

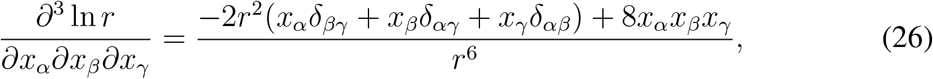

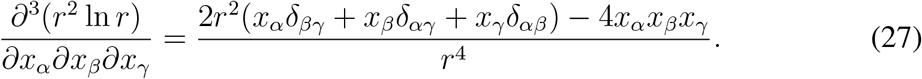

Substituting these identities directly into Eq. (24) and simplifying allows us to re-write the velocity in the following form

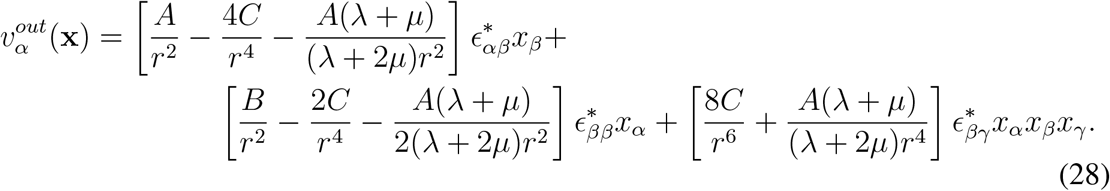

The remaining constants can be computed by enforcing continuity of the velocity field at the boundary of the inclusion where *r* = *a*. First, we know that the third term that is cubic in **x** must vanish when *r* = *a* since the velocity within the inclusion is linear in **x**.

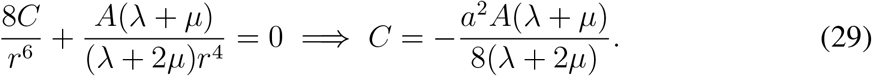

The dependence of **v**^*out*^(**x**) on *C* can now be removed yielding

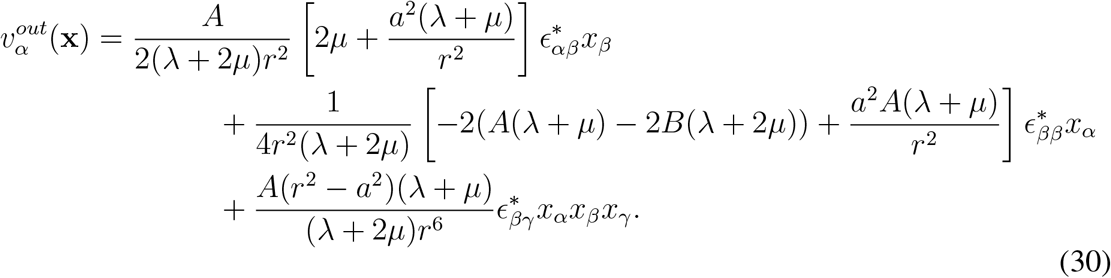

Referencing Eq. (15), we continue to enforce continuity at the boundary of the inclusion and match the coefficients of the velocities inside and outside the inclusion term by term. The term proportional to *ϵ*^*∗*^*x*_*β*_ tells us

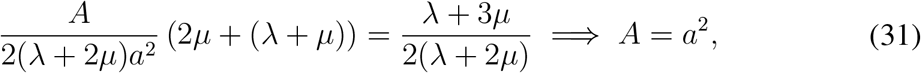

and the term proportional to 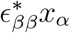 tells us

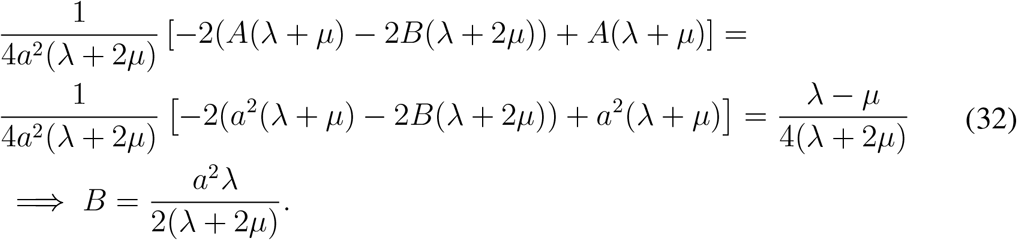

Finally we arrive at the following form for the velocity outside of the inclusion

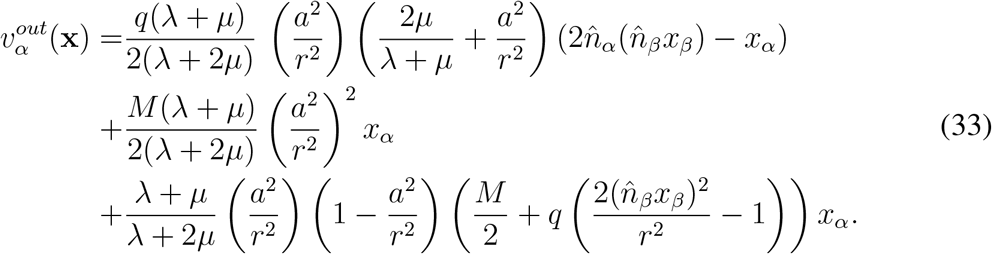

In summary, the instantaneous velocity induced by a cell division is has the following vectorial form

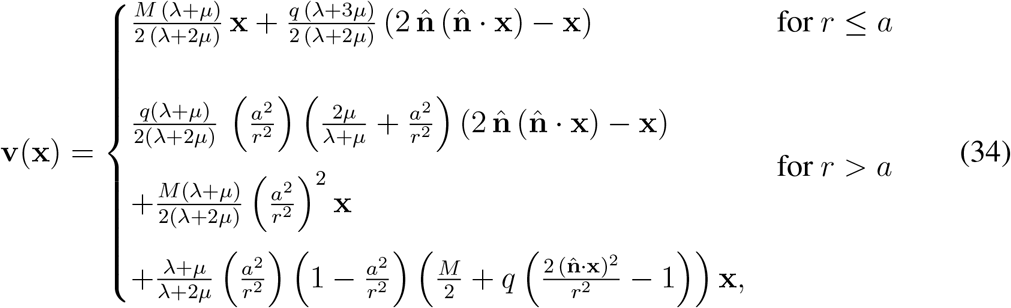

or, in terms of the effective viscosities *ν*_1_ = *τ*_*R*_ *µ* and *ν*_2_ = *τ*_*R*_ (*λ* + *µ*)

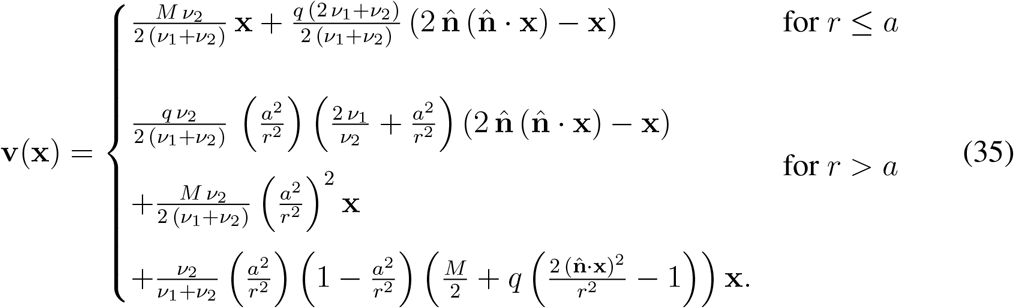

This equation can be non-dimensionalized by introducing the dimensionless parameter

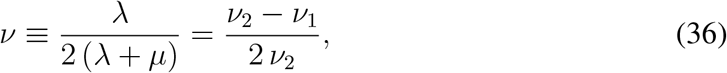

in terms of which the division velocity is

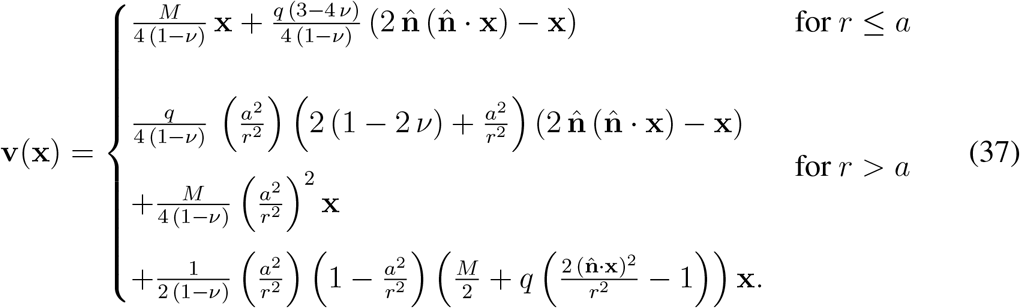

**Fig. S1.**
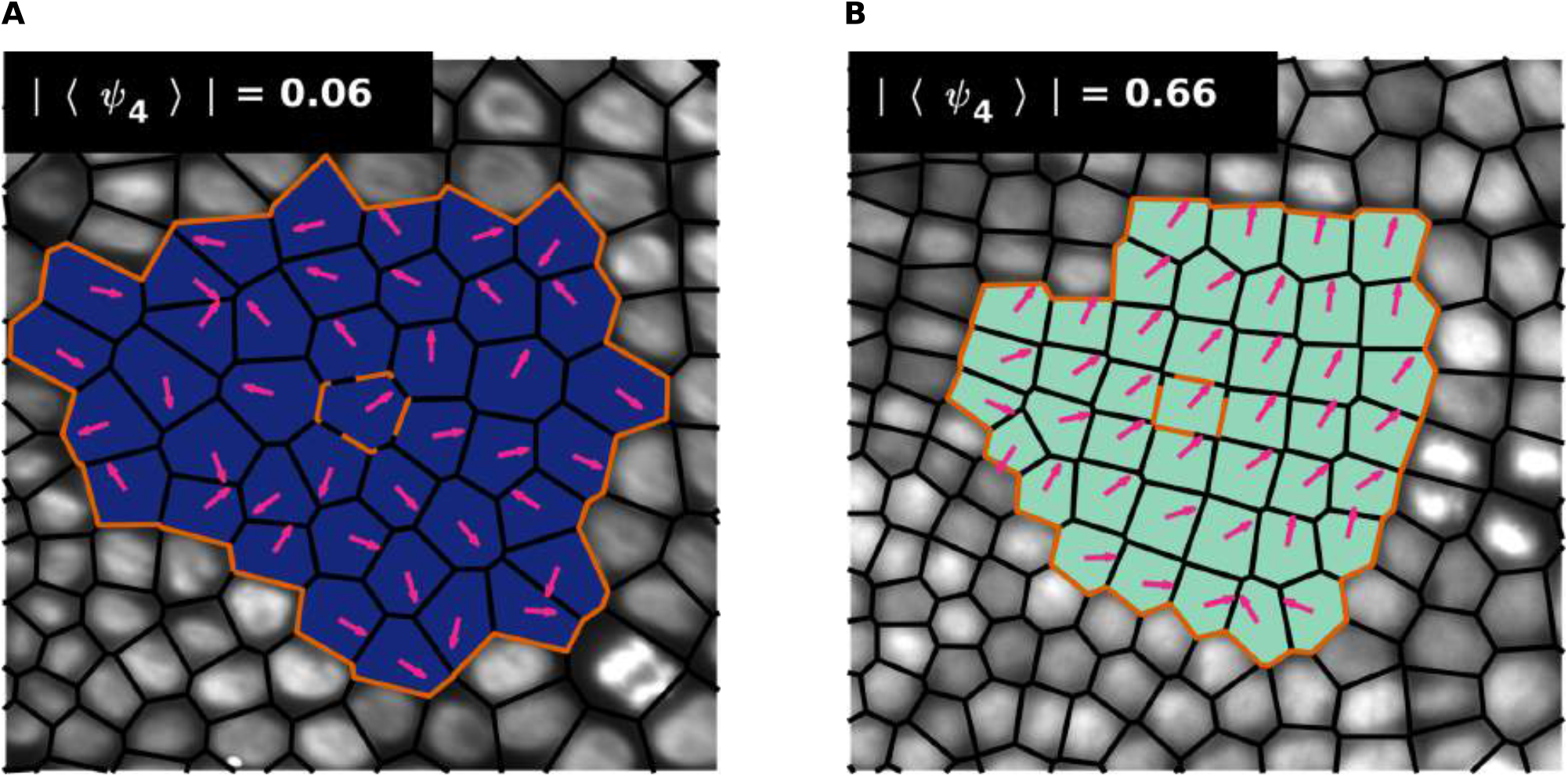
Effects of averaging the discrete fourfold orientational order parameter. (**A** and **B**) Average fourfold order parameter over the third order natural neighborhood of the single cells shown in (Fig. 1, G and H). The dotted orange boundary denotes the seed cell. The solid orange boundary denotes the averaging region. Arrows indicate the orientation of the single cell order parameter for each cell in the averaging region. (A) Average order parameter in a disordered region. (B) Average order parameter in a highly fourfold ordered region. In general, small differences in orientation will result in significantly lower mean order parameters compared to the magnitude of the single cell order parameters. This effect is especially noticeable in the high order case.

**Fig. S2.**
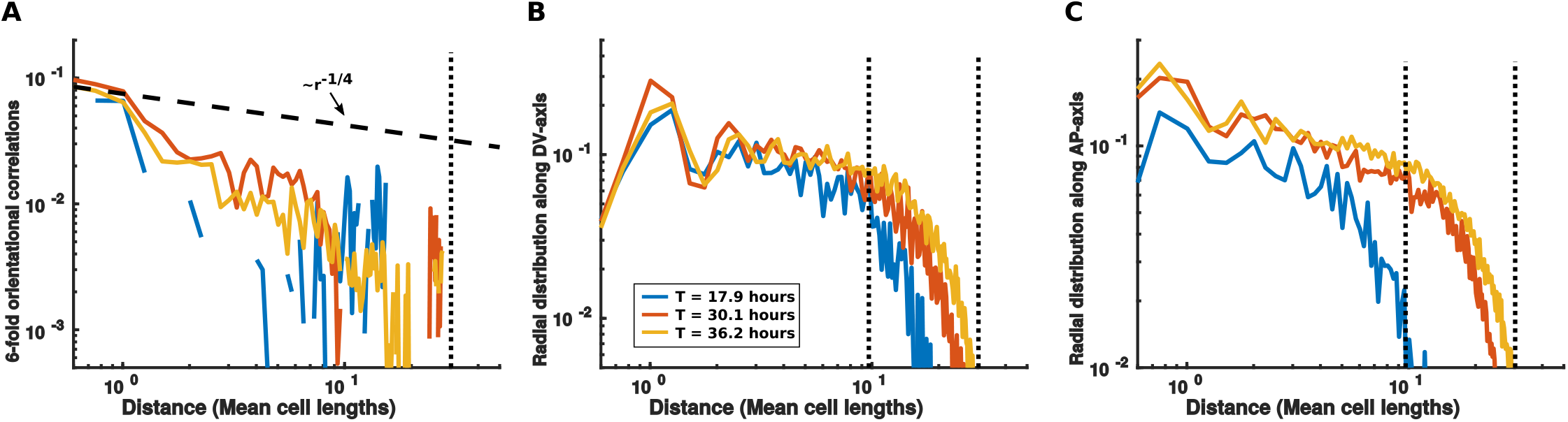
Trunk ectodermal germband exhibits neither sixfold nor translational order. (**A**) The two-point sixfold orientational correlation functions at three representative time points. All timepoint exhibit exponential decay. Vertical dotted line indicates largest lateral length scale in the system. (**B**) The pairwise radial distribution function measured along the D-V axis. Vertical dotted lines indicate length scale of the system along the A-P and D-V axes. (**C**) The pairwise radial distribution function measured along the A-P axis.

**Fig. S3.**
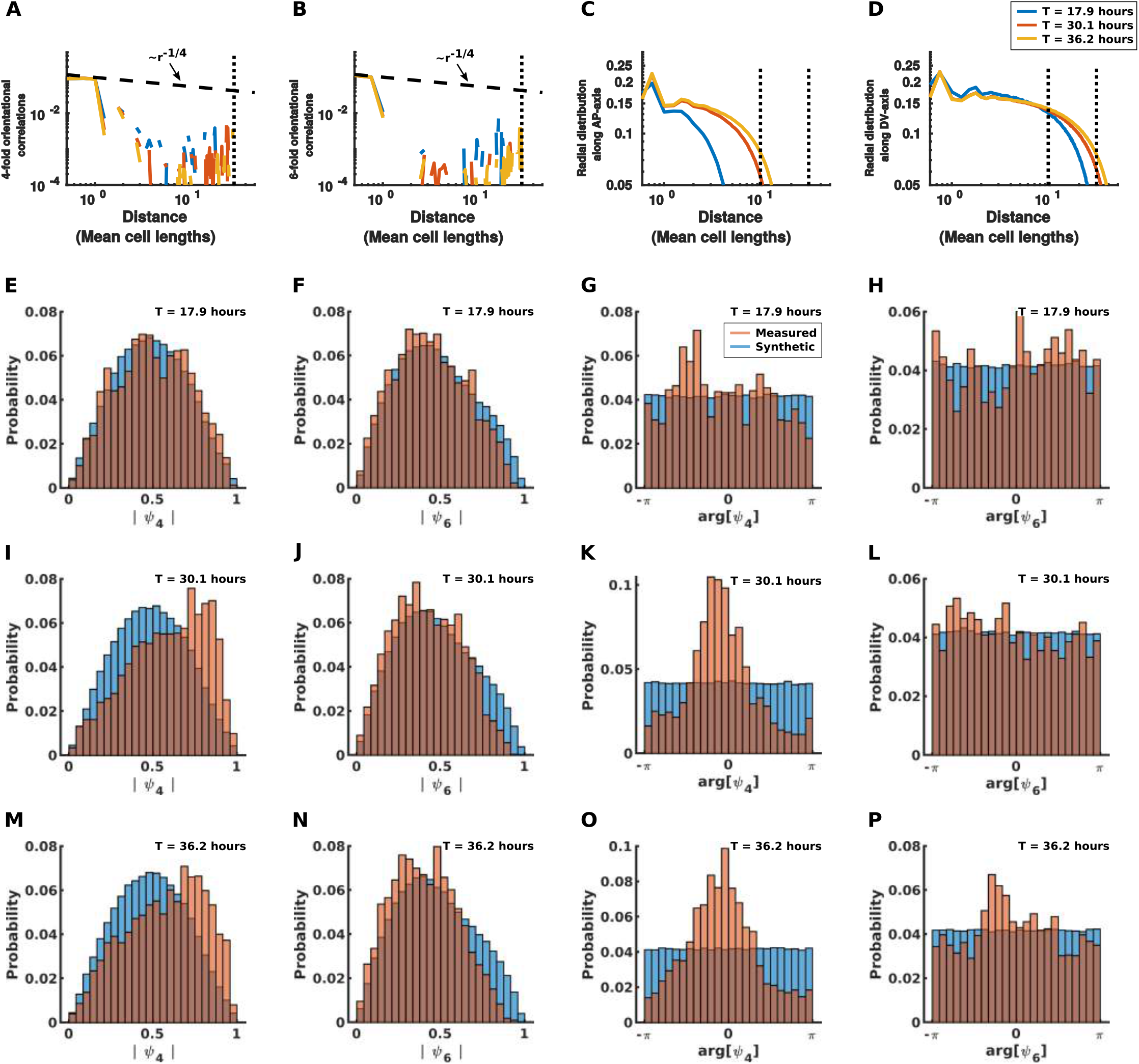
Observed order in the embryo is significant compared to disordered synthetic data sets. (**A** to **D**) Correlation functions for synthetic datas sets with the same system sizes and cell densities as the representative time points in Fig. 1J and fig. S2. (A) The two-point fourfold orientational correlation function. (B) The two-point sixfold orientational correlation function. (B) The pairwise radial distribution function measured along the A-P axis of the synthetic data set. (C) The pairwise radial distribution function measured along the D-V axis of the synthetic data set. (**E** to **H**) Comparison of the fourfold and sixfold order parameters of the measured and synthetic data sets at early times. (E) The magnitude of the fourfold order parameter. (F) The magnitude of the sixfold order parameter. (G) The phase of the fourfold order parameter. (H) The phase of the sixfold order parameter. (**I** to **L**) Comparison of the fourfold and sixfold order parameters of the measured and synthetic data sets at intermediate times. (I) The magnitude of the fourfold order parameter. (J) The magnitude of the sixfold order parameter. (K) The phase of the fourfold order parameter. (L) The phase of the sixfold order parameter. (**M** to **P**) Comparison of the fourfold and sixfold order parameters of the measured and synthetic data sets at late times. (M) The magnitude of the fourfold order parameter. (N) The magnitude of the sixfold order parameter. (O) The phase of the fourfold order parameter. (P) The phase of the sixfold order parameter.

**Fig. S4.**
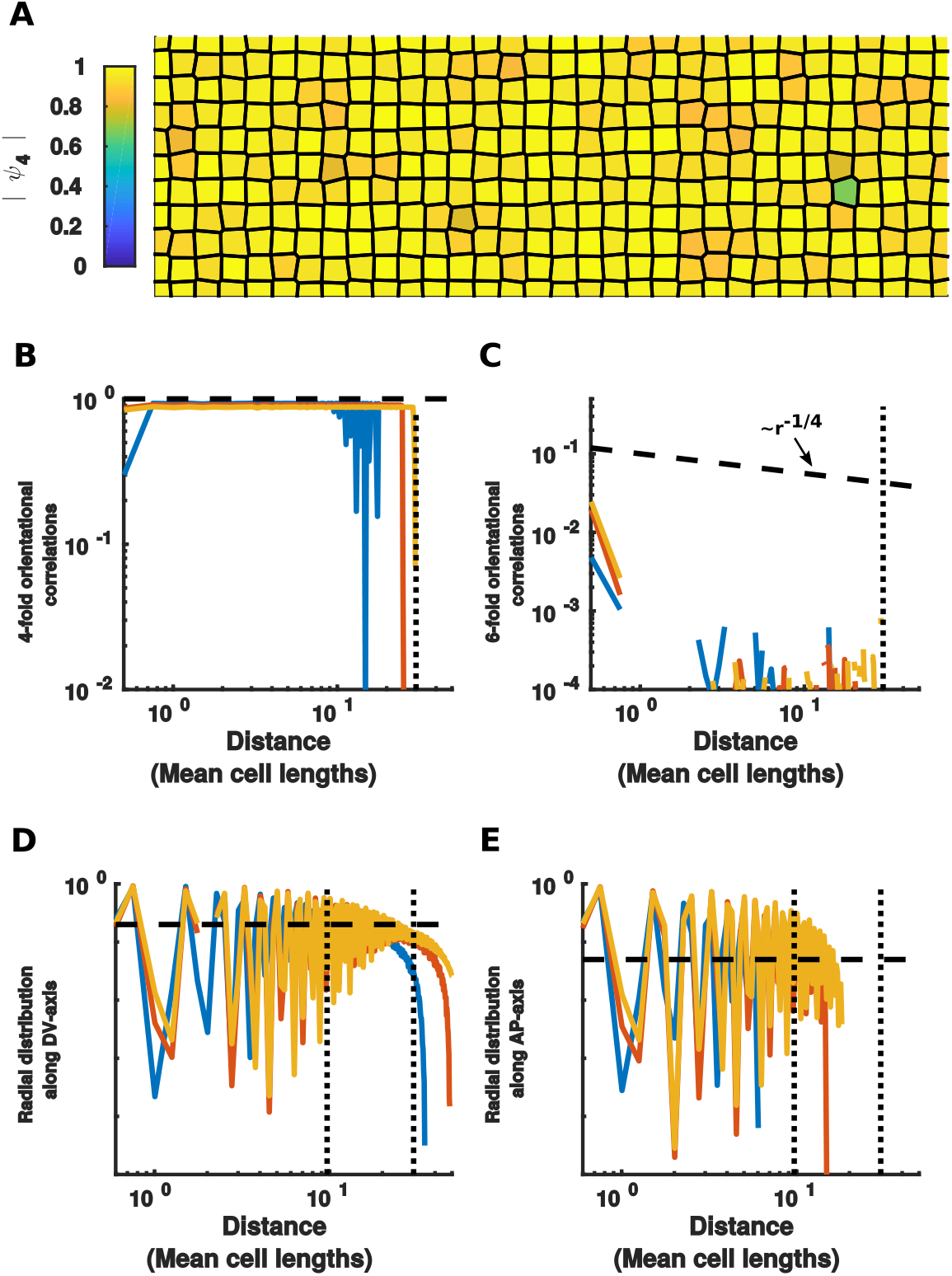
The germband does not exhibit translational order. (**A**) An example translationally ordered synthetic data set with the same spatial dimensions and cell density as the germband at *T* = 36.2 hours. Color of cells indicates the magnitude of the fourfold orientational order parameters. (**B** to **E**) Correlation functions for synthetic data sets with the same system sizes and cell densities as the representative time points in Fig. 1J and fig. S2. (B) The two-point fourfold orientational correlation function. (C) The two-point sixfold orientational correlation function. (D) The pairwise radial distribution function measured along the A-P axis of the synthetic data set. The horizontal dashed line is a guide for the eye. (E) The pairwise radial distribution function measured along the D-V axis of the synthetic data set.

**Fig. S5.**
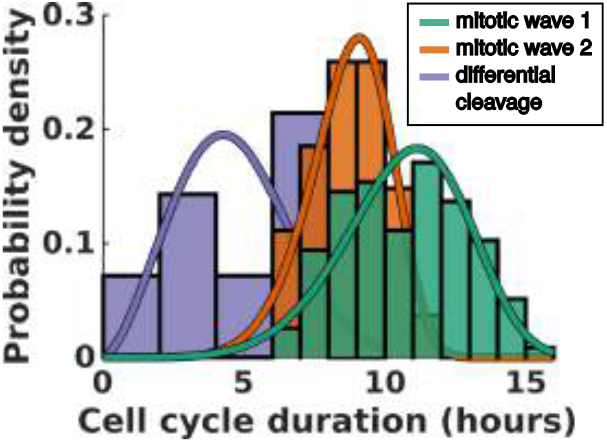
Distribution of cell cycle durations throughout growth. Solid lines are Weibull distributions fit to the cell cycle duration histograms. Average duration and standard deviation for mitotic wave 1 is 10.7 ± 2.2 hours from *N* = 117 cell cycles. Average duration and standard deviation for mitotic wave 2 is 8.7 ± 1.5 hours from *N* = 27 cell cycles. Average duration and standard deviation for differential cleaveage is 4.6 ± 2.0 hours from *N* = 7 cell cycles.

**Fig. S6.**
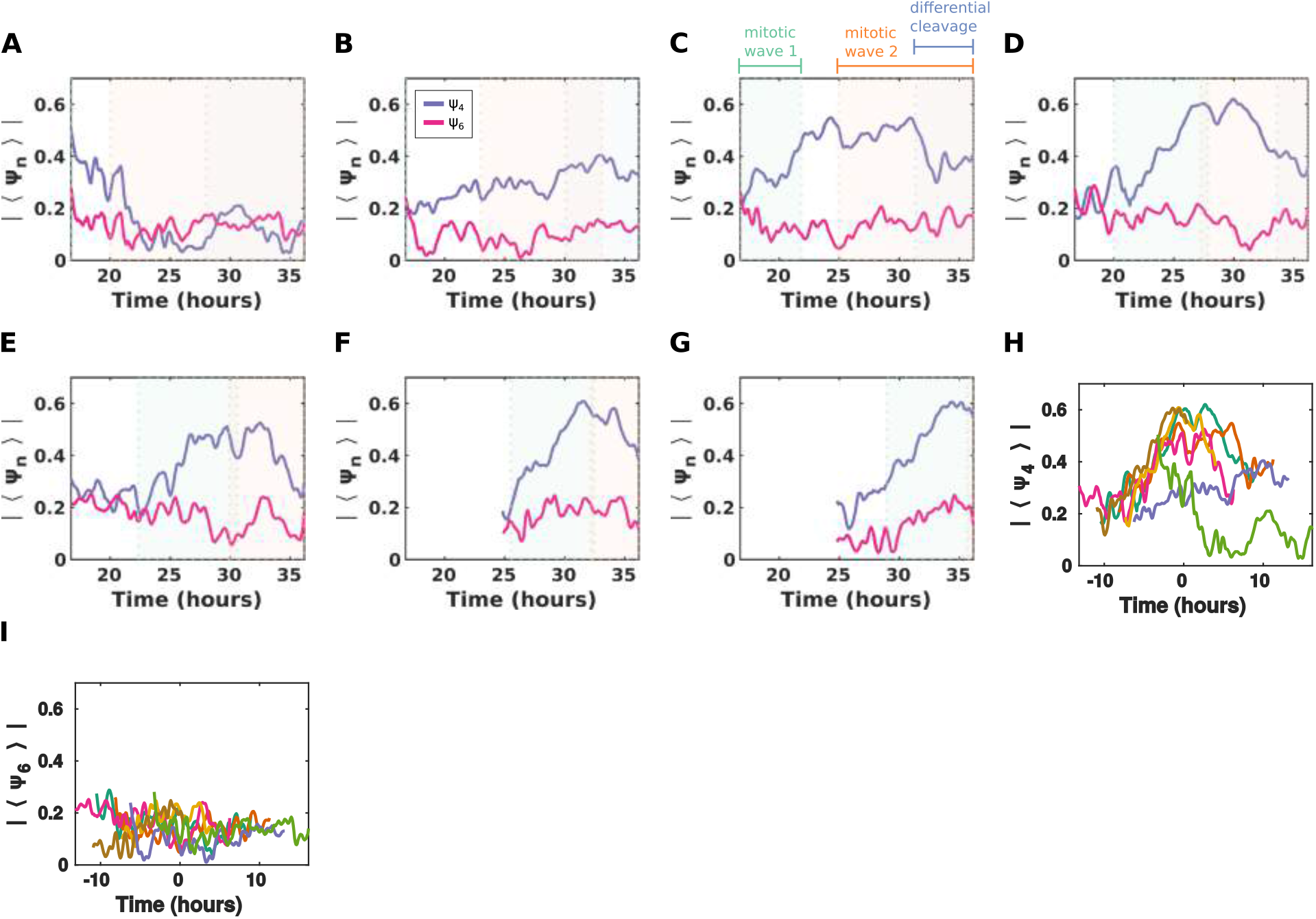
Temporal profile of orientational order is similar across different parasegments. (**A** to **G**) Per-parasegment profile of of the magnitude of the average fourfold and sixfold orientational order parameters over time. Ordering of panels (A to G) mirrors anterior to posterior ordering of the physical segments (i.e. the parasegment shown in (A) is anterior to the parasegment shown in (B), etc.). (**H**) The magnitude of the average four-fold orientational order parameter for each parasegment shifted in time so that *T* = 0 corresponds to the first cell division of mitotic wave 2. (**I**) The magnitude of the average sixfold orientational order parameter for each parasegment shifted in time so that *T* = 0 corresponds to the first cell division of mitotic wave 2.

**Fig. S7.**
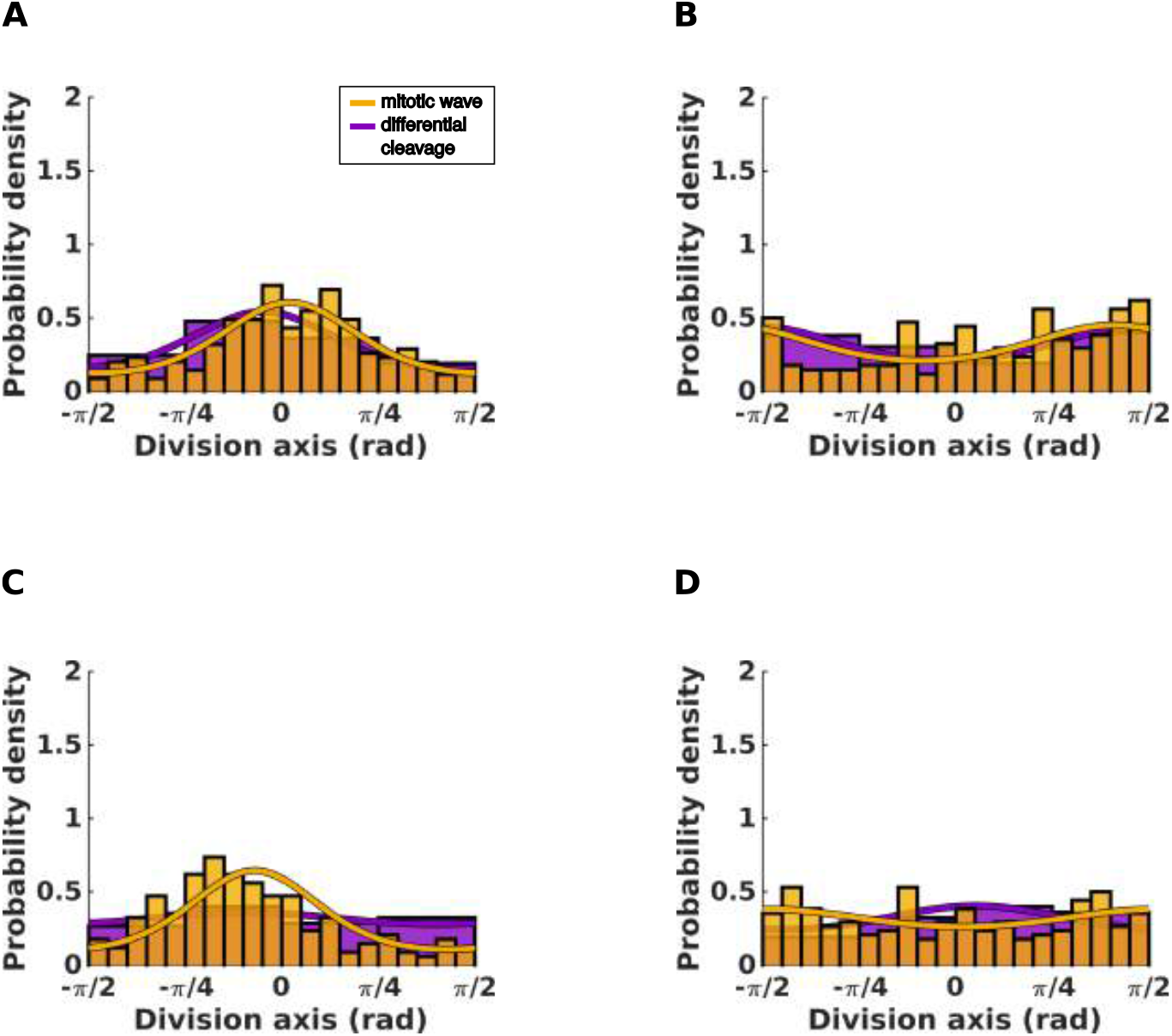
Orientations of cell divisions are not predicted by local mechanical or geometric signals. (**A**) Histogram of orientations of cell divisions relative to the principal axis of the strain-rate of the corresponding parent cell integrated over the hour preceding cell division. (**B**) Histogram of the orientations of cell divisions relative to the axis of elongation of the corresponding parent cell averaged over the hour preceding cell division. (**C**) Histogram of the orientations of cell divisions relative of the orientation of the fourfold order parameter of the corresponding parent cell averaged over the hour preceding cell division. (**D**) Histogram of the orientations of cell divisions relative to the orientation of the sixfold order parameter of the corresponding parent cell averaged over the hour preceding cell division.

**Fig. S8.**
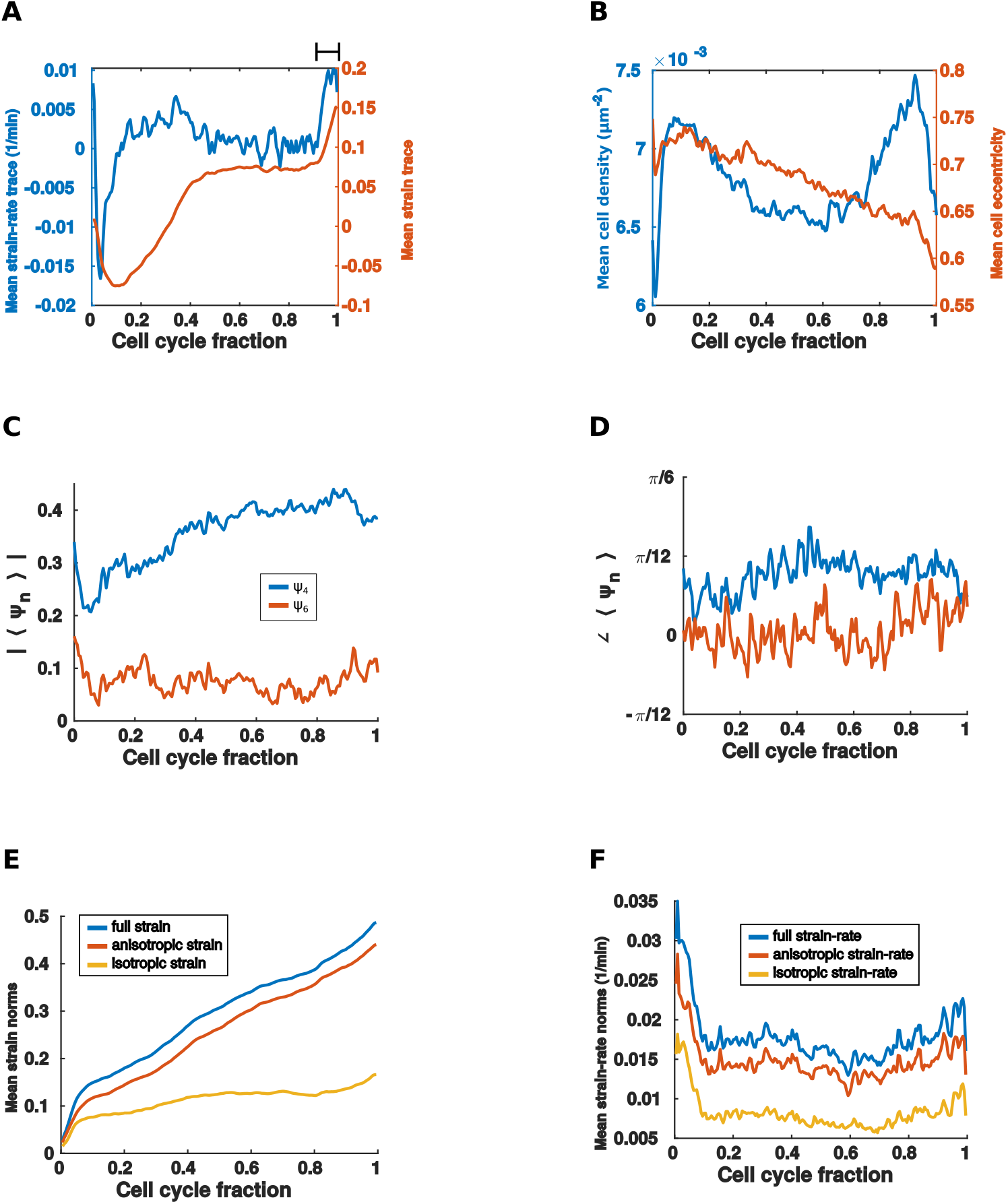
Timing of cell divisions are not predicted by local mechanical or geometric signals. (**A** to **F**) Various geometric and mechanical fields averaged over a normalized cell cycle. Average includes cell cycles from both mitotic waves and differential cleavage. (A) The trace of the mean strain-rate tensor (left) and the trace of the mean cumulative strain tensor (right). The black bar in the upper right hand corner indicates the growth phase preceding cell division, i.e. cell division has already been initiated in this phase. (B) Mean cell density (left) and mean eccentricity of an ellipse fit to each cell (right). (C) The magnitude of the average fourfold and sixfold order parameters. (D) The orientation of the average fourfold and sixfold order parameters relative to the A-P axis. (E) The Frobenius norm of the average full cumulative strain tensor, the anistropic part of the the average strain tensor, and the isotropic part of the average strain tensor. (F) The Frobenius norm of the full average strain-rate tensor, the anisotropic part of the average strain-rate tensor, and the isotropic part of the average strain-rate tensor.

**Fig. S9.**
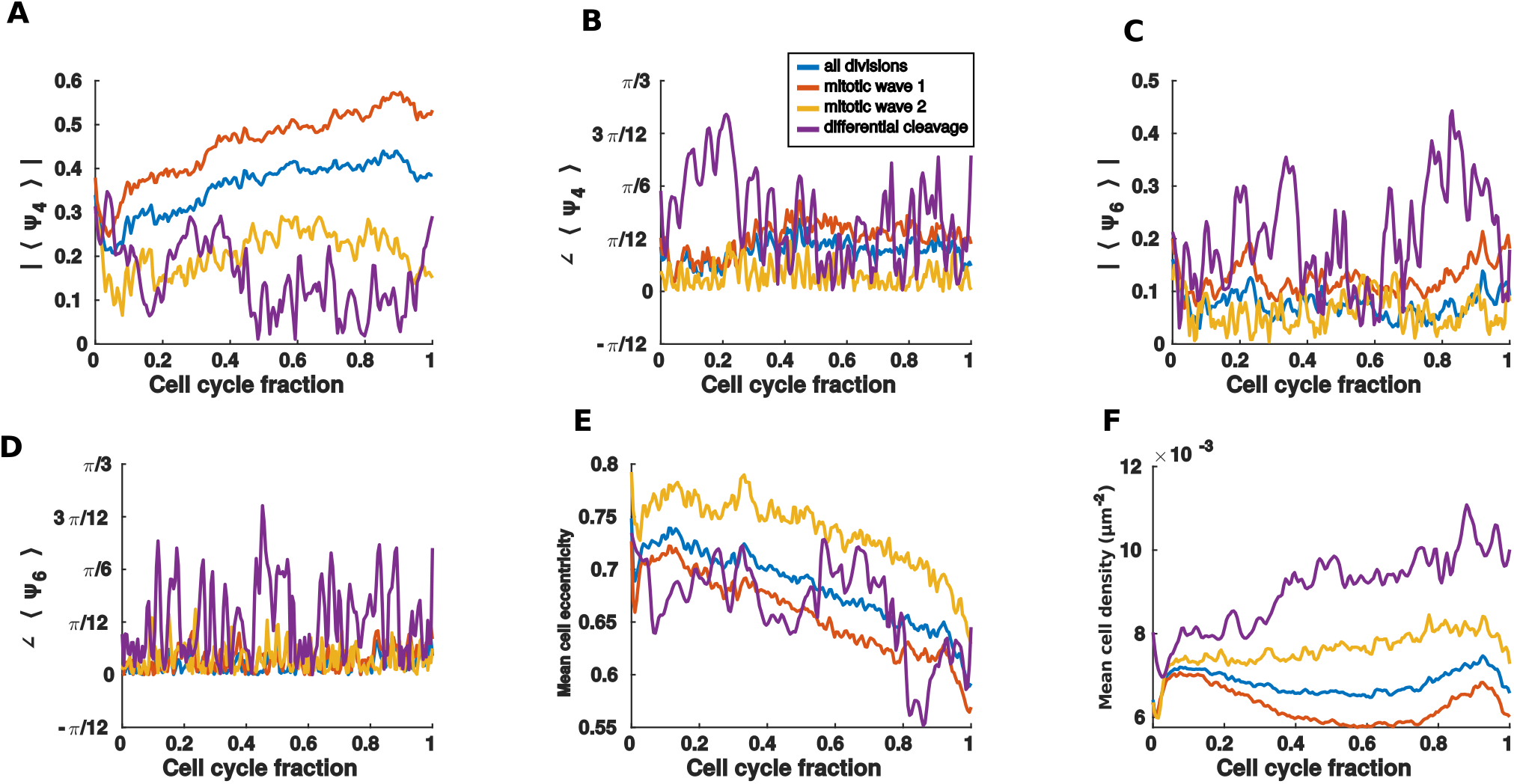
Timing of cell divisions are not predicted by local geometric signals (cont.). (**A** to **F**) Various geometric fields averaged over a normalized cell cycle. Separate averages are included for each mitotic wave and differential cleavage. (A) The magnitude of the mean fourfold orientational order parmeter. (B) The orientation of the mean fourfold orientational order parmeter. (C) The magnitude of the mean sixfold orientational order parameter. (D) The orietnation of the mean sixfold orientational order parameter. (E) The mean eccentricity of an ellipse fit to each cell. (F) The mean cell density.

**Fig. S10.**
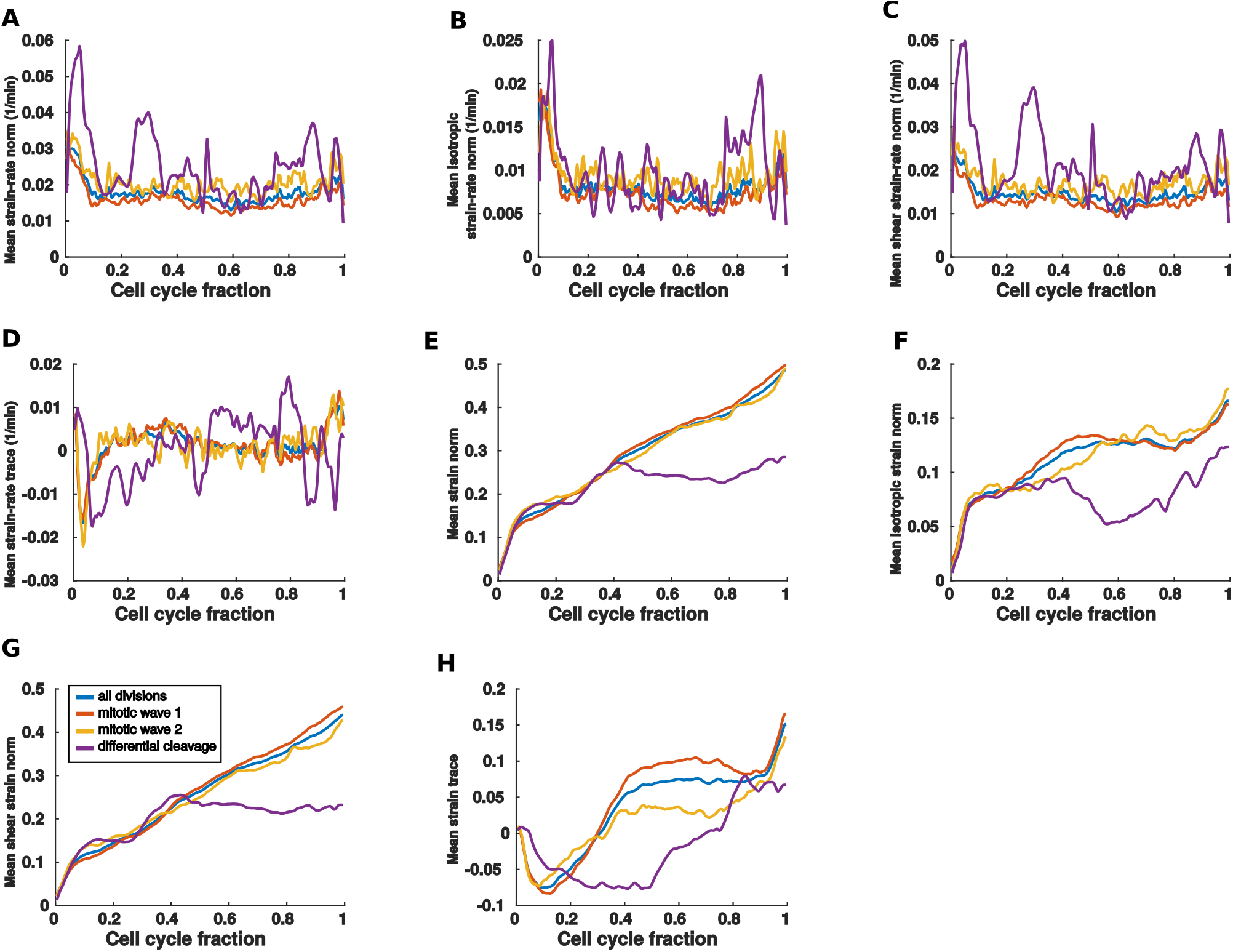
Timing of cell divisions are not predicted by local mechanical signals (cont.). (**A** to **H**) Various mechanical fields averaged over a normalized cell cycle. Separate averages are included for each mitotic wave and differential cleavage. (A) The Frobenius norm of the mean strain-rate tensor. (B) The Frobenius norm of the isotropic part of the mean strain-rate tensor. (C) The Frobenius of the anisotropic part of the mean strain-rate tensor. (D) The trace of the mean strain-rate tensor. (E) The Frobenius norm of the mean cumulative strain tensor. (F) The Frobenius norm of the isotropic part of the mean cumulative strain tensor. (G) The Frobenius norm of the anisotropic part of the mean cumulative strain tensor. (H) The trace of the mean cumulative strain tensor.

**Fig. S11.**
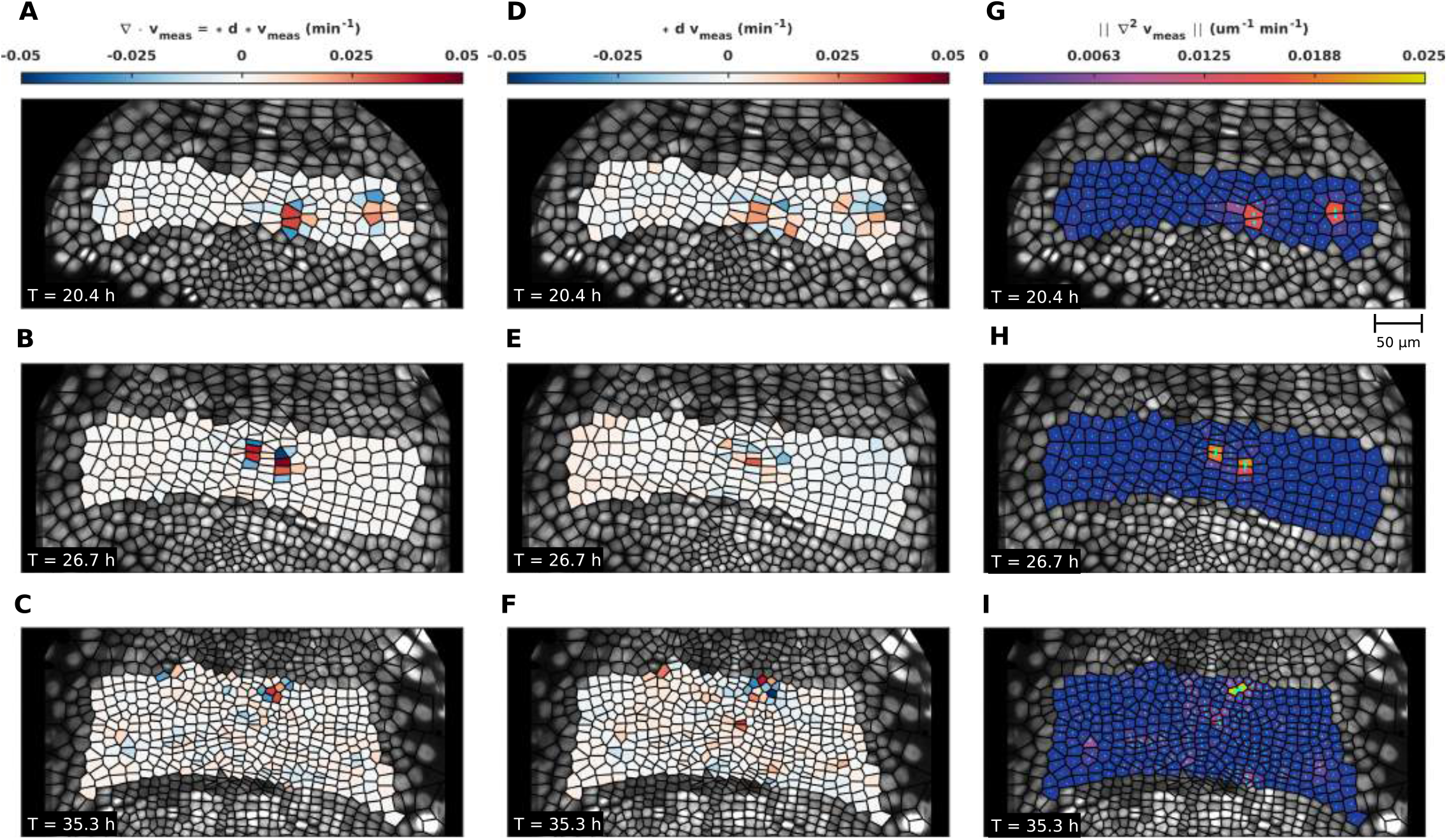
Gradients of the measured tissue velocity fields. (**A** to **C**) The divergence of the measured tissue velocity fields for three representative time points. Explicit construction in term of exterior derivatives and Hodge stars is shown in (A). (**D** to **F**) The ‘curl’ of the measured tissue velocity fields for three representative time point. Explicit construction in terms of exterior derivatives and Hodge stars is shown in (D). (**G** to **I**) The Laplacian of the measured tissue velocity fields for three representative time points.

**Fig. S12.**
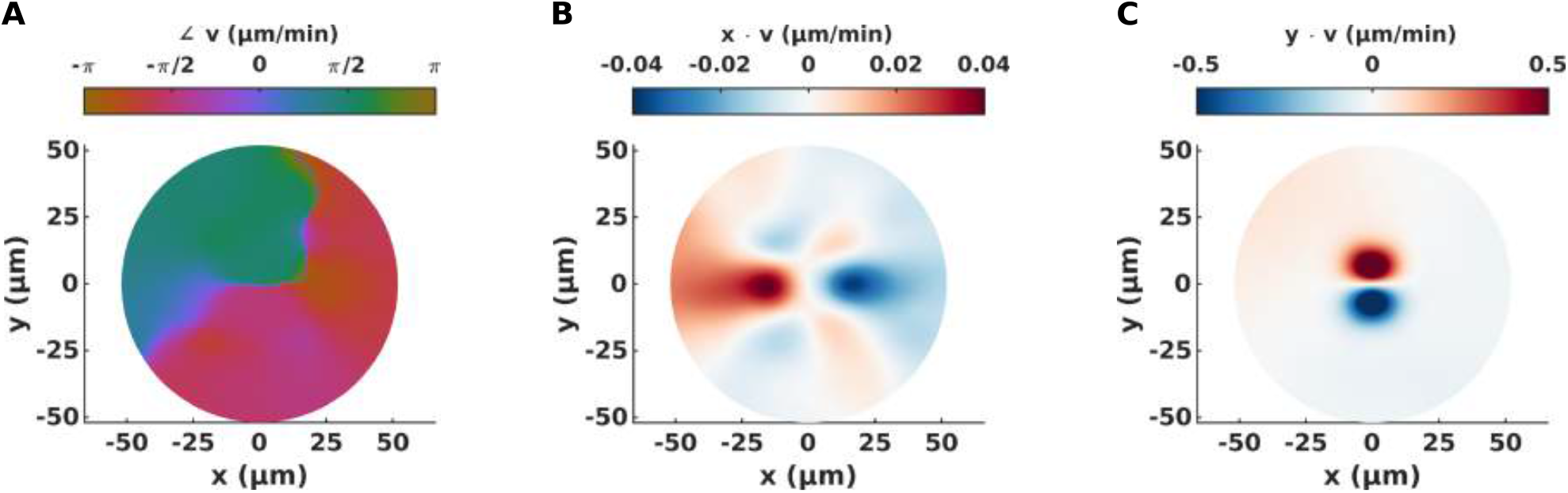
The velocity of the mean cell division event. (**A**) The orientation of the velocity of the mean cell division event. (**B**) The *x*-component of the velocity of the mean cell division event. (**C**) The *y*-component of the velocity of the mean cell division event.

**Fig. S13.**
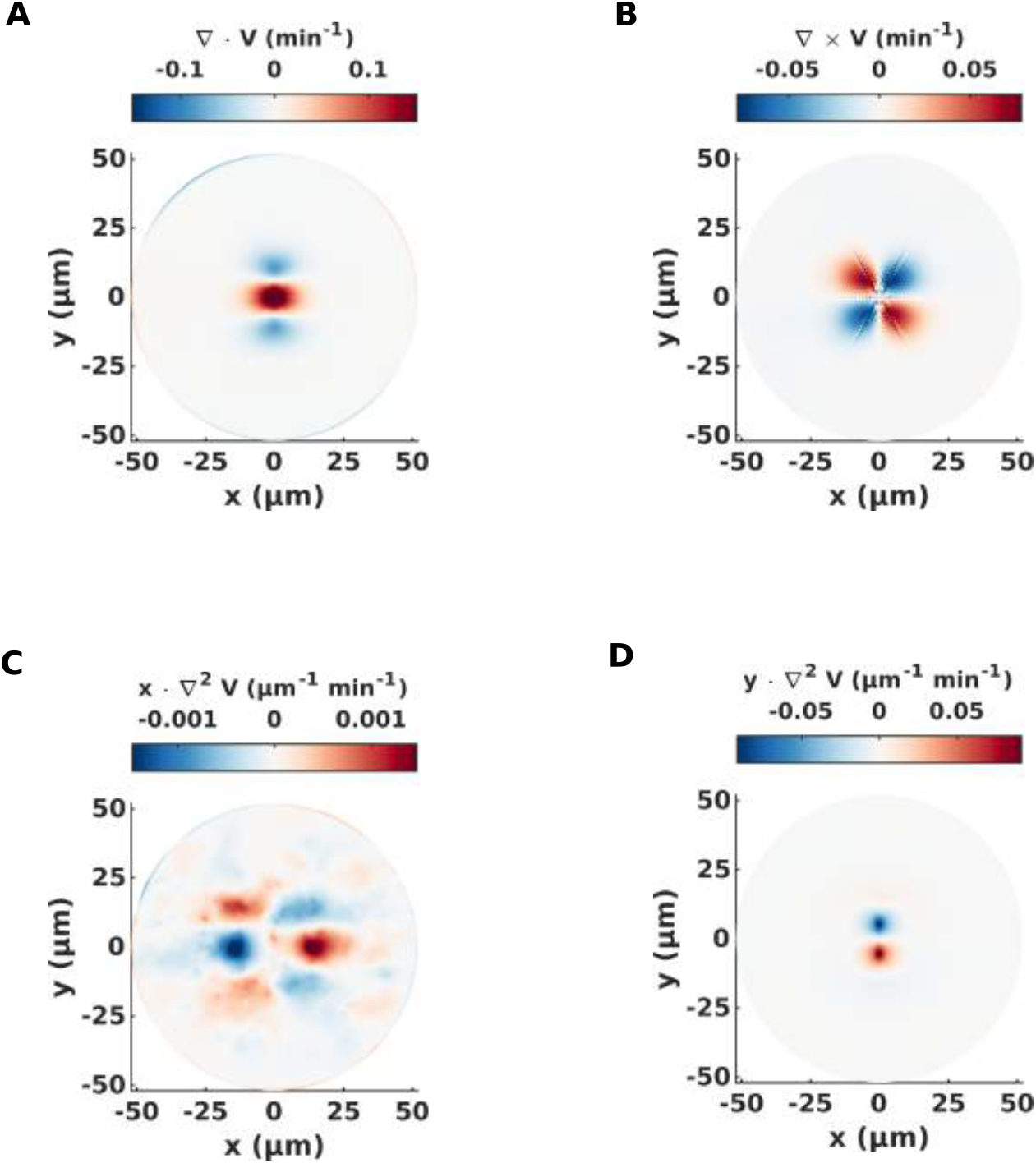
Gradients of the velocity of the mean cell division event. (**A**) The divergence of the mean cell division velocity. (**B**) The ‘curl’ of the mean cell division velocity. (**C**) The *x*-component of the Laplacian of the mean cell division velocity. (**D**) The *y*-component of the Laplacian of the mean cell division velocity.

**Fig. S14.**
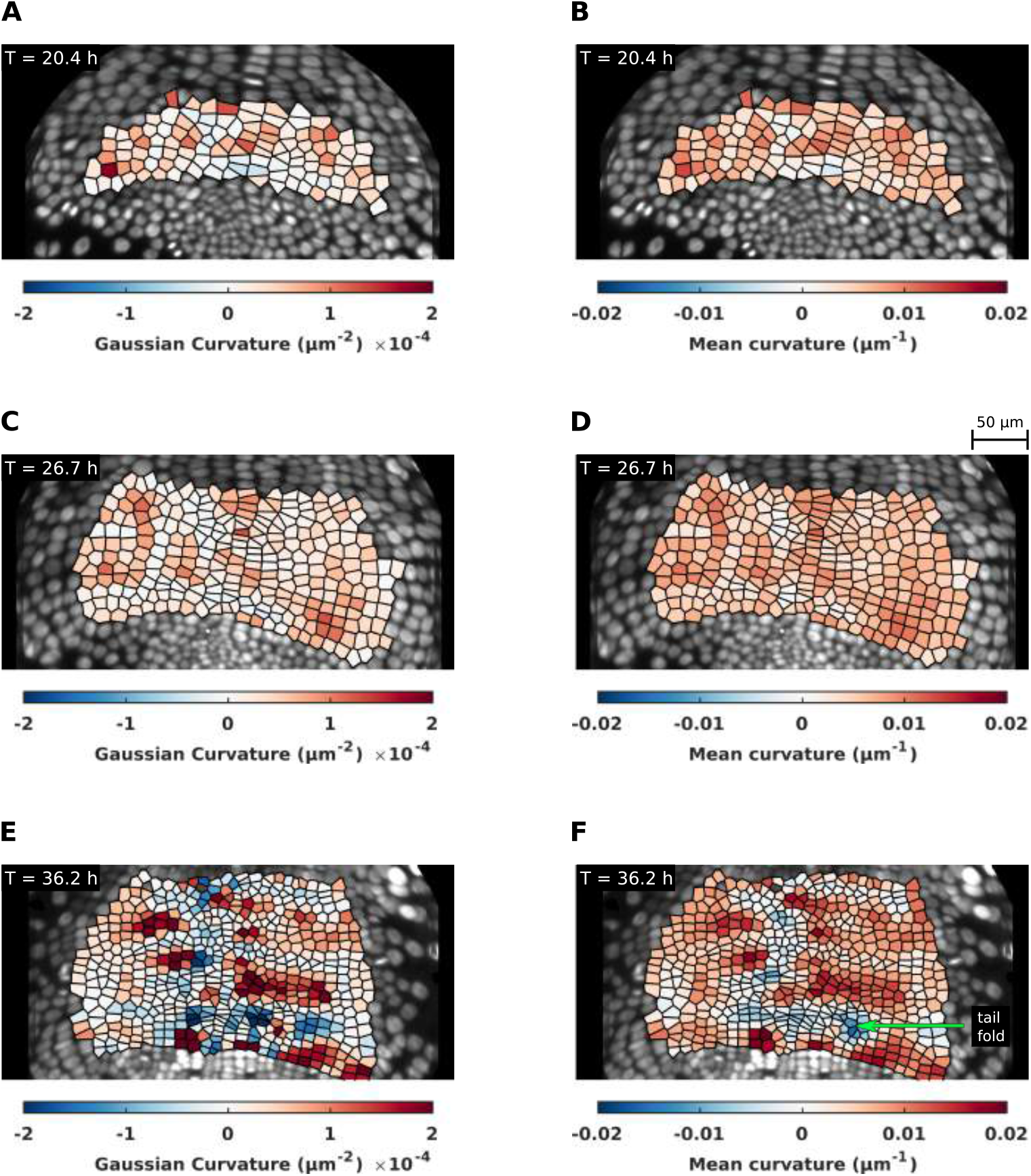
Surface curvatures in the growing *Parhyale* embryo. (**A**) Per-cell Gaussian curvature at early times. (**B**) Per-cell mean curvature at early times. (**C**) Per-cell Gaussian curvature at intermediate times. (**D**) Per-cell mean curvature at intermediate times. (**E**) Per-cell Gaussian curvature at late times. (**F**) Per-cell mean curvature at late times.

**Fig. S15.**
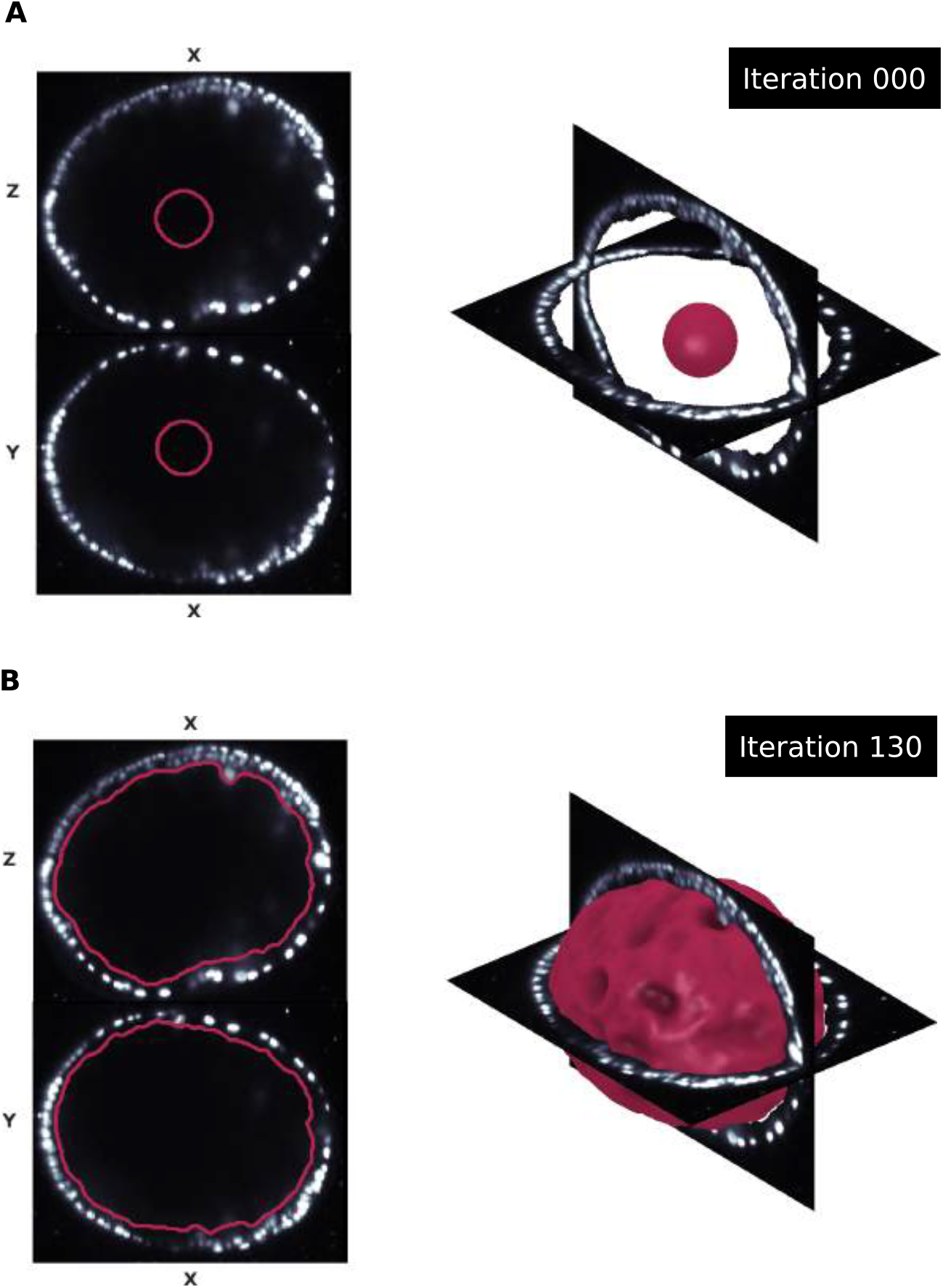
Surface extraction via level sets. An illustration of the morphological snakes method used to extract the surface of interest in the growing embryo. (**A**) The spherical initial condition (zero completed iterations). (**B**) The segmented surface.

**Fig. S16.**
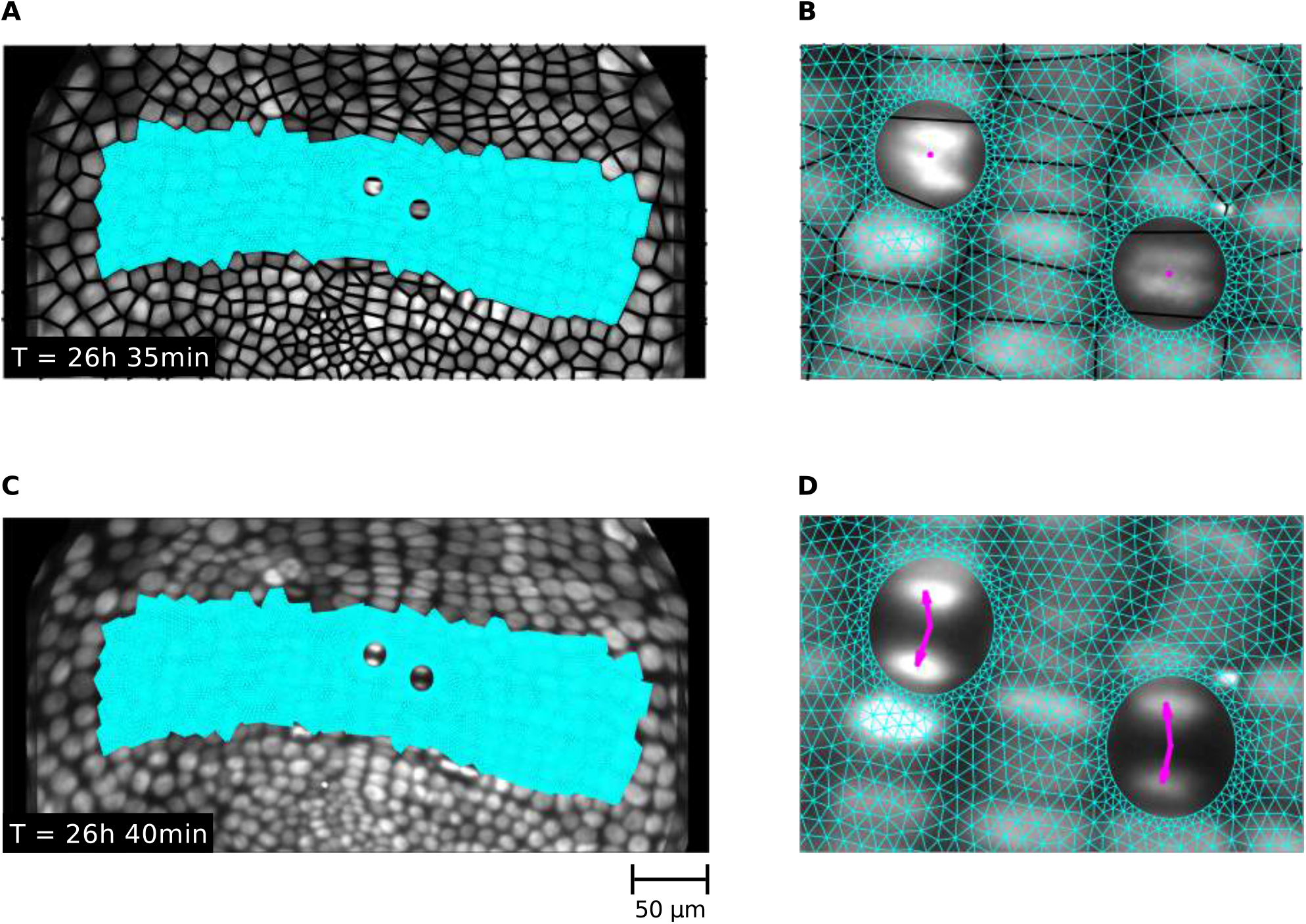
Numerical solution of the fluid mechanical equations of motion. Examples of the triangular mesh used to implement a finite element method (FEM) solution to the equations of motion predicting tissue velocities from cell divisions. (**A**) The undeformed mesh triangulation. Circular holes have been removed surrounding cells about to divide. **(B)** A zoomed-in visualization of the undeformed mesh near the cells about to divide. (**C**) The deformed mesh at the subsequent time. Mesh vertex velocities have been scaled by a constant pre-factor (x5) to improve visibility. (**D**) A zoomed-in visualization of the mesh near the recently divided cells. Note that the initially circular holes are deformed into ellipses.

**Table S1.**
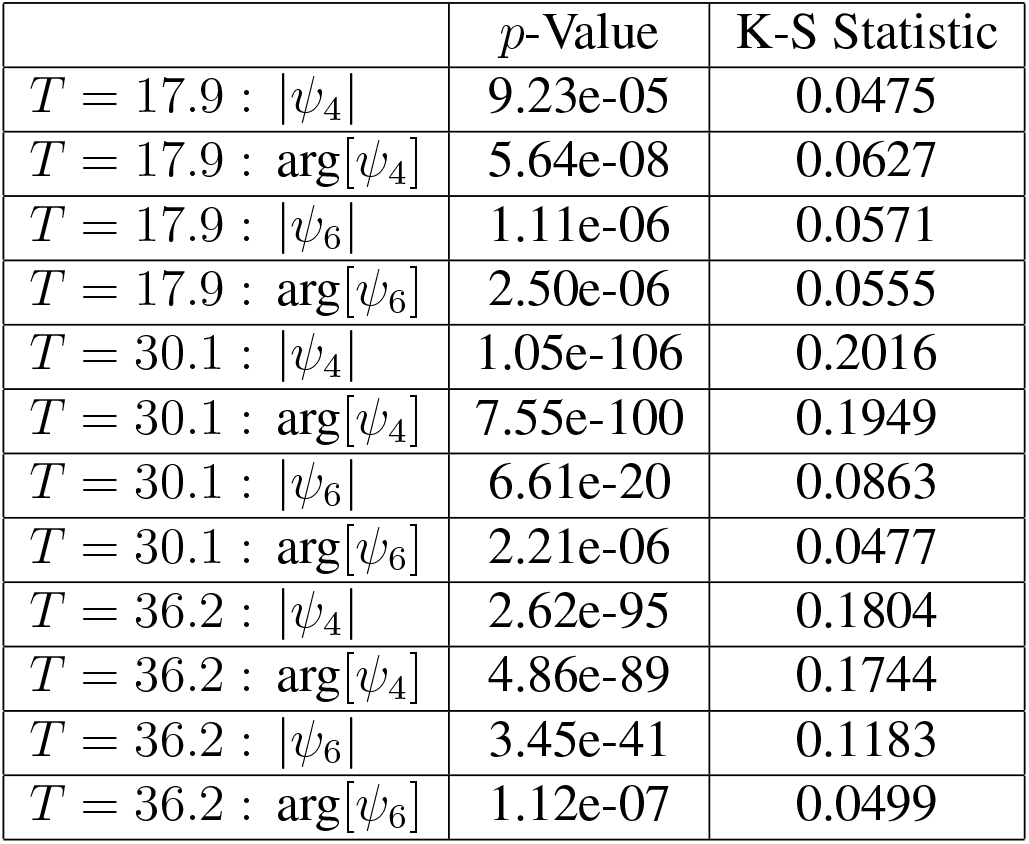
Statistical comparison of orientational order in measured and synthetic data sets. The Kolmogorov-Smirnov (K-S) statistic and asymptotic *p*-values for the distributions shown in fig. S3. Details on calculation in materials and methods.

**Movie S1. Surface extraction and pullback construction** The first segment of the movie illustrates the morphological snakes method for surface detection. Left panels show cross sections of the data and the expanding interior region defined by the morphological snakes algorithm. The right panel shows the same process in a 3D view. The next segment of the movie visualizes how a disk-like region of interest can be extracted from a sphere-like surface and mapped into the plane.

**Movie S2. 3D dynamic surface growth visualization** The trunk ectodermal germband of a growing *Parhyale* embryo shown as a 3D dynamic surface.

**Movie S3. 2D conformal germband growth visualization.** The same trunk ectodermal germband region shown in Movie S2 pulled back conformally to the unit disk and visualized in 2D.

**Movie S4. 2D whole embryo growth visualization.** An ‘As-Rigid-As-Possible’ pull-back of the entire embryo in 2D. Includes the head ectoderm in addition to the trunk ectodermal germband.

**Movie S5. Oriented cell division analysis.** The oriented division of a single cell along the A-P axis. The magenta line shows the dynamic orientation of the A-P axis. The cyan line shows the orientation of the cell division. Prior to cell division, the orientation of the nuclei is found by fitting an ellipse to the corresponding dynamic nuclear segmentation and extracting the minor axis of that fit ellipse. Following cell division, the orientation is taken to be the line separating daughter cell centroids.

**Movie S6. Division choreography of a single parasegment pre-cursor row (PSPR).** Shows the characteristic proliferation of a single PSPR into two rows, and subsequently into four rows. A single cell is outlined to highlight this choreography on a smaller scale.

**Movie S7. Division choreography of a single cell.** A close-up of the single highlighted cell lineage in Movie S6 playing out the same division choreography.

